# HTRA1 deficiency in *COL4A1* mutant hiPSC-derived astrocytes: a convergent mechanism of cerebral small vessel disease

**DOI:** 10.64898/2026.05.12.724691

**Authors:** Xuewei Qi, Alessandra Granata, Tom Van Agtmael, Sanjay Sinha, M. Zameel Cader, Hugh S Markus, Stuart M Allan, Karen Horsburgh, Tao Wang

**Affiliations:** Division of Evolution, Infection and Genomics, School of Biological Sciences, Faculty of Biology, Medicine and Health, The University of Manchester, Manchester M13 9PT, UK; Stroke Research Group, Department of Clinical Neurosciences Department, University of Cambridge, Cambridge CB2 0QQ, UK; School of Cardiovascular and Metabolic Health, University of Glasgow, Glasgow G12 8QQ, UK; Cambridge Stem Cell Institute, Jeffrey Cheah Biomedical Centre, Cambridge CB2 0AW, UK; Translational Molecular Neuroscience Group, Weatherall Institute of Molecular Medicine, Nuffield Department of Clinical Neurosciences, University of Oxford, Oxford OX3 9DU, UK; Division of Neuroscience, School of Biological Sciences, Faculty of Biology, Medicine and Health, The University of Manchester, Manchester M13 9PT, UK; Geoffrey Jefferson Brain Research Centre, Manchester Academic Health Science Centre, Northern Care Alliance NHS Foundation Trust, The University of Manchester, Salford M6 8HD, UK; Institute for Neuroscience and Cardiovascular Research, University of Edinburgh, Edinburgh EH16 4SB, UK

**Keywords:** *COL4A1*, Cerebral small vessel disease, *HTRA1*, extracellular matrix, blood-brain barrier, human induced pluripotent stem cells

## Abstract

Cerebral small vessel disease (cSVD) is a major contributor to stroke and cognitive decline, ultimately leading to vascular dementia (VaD). Genetic factors play a key role in disease susceptibility and progression, and variants in *COL4A1* cause one of the most common forms of genetic cSVD. *COL4A1* encodes the α1 chain of collagen type IV which is the major structural component of the basement membrane, a specialised extracellular matrix (ECM) structure, in the vasculature. In addition to this vascular basement membrane (vBM), in the central nervous system (CNS), the neurovascular unit (NVU) also has a unique parenchymal basement membrane (pBM) that is largely produced by astrocytes and forms a critical interface anchoring astrocyte end-feet to vascular cells. Together, these BMs play essential roles in regulating blood-brain barrier (BBB) function. However, the role of the pBM in cSVD has been relatively less investigated compared to the vBM. The lack of relevant human disease models that faithfully recapitulate the pBM, makes it difficult to dissect pBM-related cell-cell and cell-matrix interactions specific to cSVD, hindering the identification of effective therapeutic targets. In this study, we hypothesised that astrocyte-mediated ECM remodelling contributes to BBB dysfunction in *COL4A1*-associated cSVD. To investigate this, human induced pluripotent stem cells (iPSCs) derived from a patient carrying the *COL4A1^G^*^755^*^R^*variant and its isogenic control line were differentiated into astrocytes and brain microvascular endothelial cells (BMECs). Comparing to isogenic controls, the *COL4A1^G^*^755^*^R^*astrocytes significantly reduced the expression of ECM-related genes and increased glutamate uptake. ECM preparations from *COL4A1^G^*^755^*^R^* astrocytes significantly damaged the tight junction (TJ) structure formed by control iPSC-BMECs and failed to rescue the TJ integrity in *COL4A1^G^*^755^*^R^* BMECs. The secretome from *COL4A1^G^*^755^*^R^*astrocytes exacerbated the ECM defects in *COL4A1^G^*^755^*^R^* BMECs. Most importantly, *COL4A1^G^*^755^*^R^* astrocytes exhibited reduced expression of HTRA1, a serine protease that regulates both ECM turnover and homeostasis, and increased TGF-β signalling. Functional rescue by recombinant human HTRA1 protein rescues TJ defects in *COL4A1^G^*^755^*^R^* BMECs, and normalized TGF-β signalling and glutamate uptake in *COL4A1^G^*^755^*^R^* astrocytes. Together, these findings define a previously unrecognised astrocyte-driven pBM mechanism in *COL4A1*-associated cSVD and highlight HTRA1 in ECM remodelling as a therapeutic target for cSVD.

## Introduction

Cerebral small vessel disease (cSVD) is the leading cause of stroke and vascular cognitive impairment, contributing to approximately 25% of all strokes and nearly 50% of dementia cases worldwide^1–3^. The burden of cSVD is expected to increase substantially in coming decades due to increased aging globally^4^. Aetiology of cSVD is heterogeneous with the two major subtypes being arteriosclerosis resulting in “sporadic” SVD, for which the major risk factors include hypertension, diabetes and smoking, and cerebral amyloid angiopathy (CAA) that is related to amyloid deposition in the more superficial cerebral vessels. Monogenic causes of cSVD are also being increasingly recognised, the most common being due to mutations in the *NOTCH3*, *HTRA1*, and *COL4A1/2* genes. Moreover, common non-coding and rare coding variants in *COL4A1* and *COL4A2* also contribute to sporadic late-onset cSVD with intracerebral haemorrhage^5, 6^. Thus, increasing evidence suggests these mutations and many of the common genetic variants associated with sporadic cSVD^7–9^, share disease mechanisms that converge on disruption of the extracellular matrix (ECM) and associated proteins comprising the matrisome^2, 10^. This raises the possibility that therapeutically targeting matrisome disruption could offer novel treatment approaches. To achieve this, more detailed understanding of the underlying molecular pathways is required.

*COL4A1* and *COL4A2* genes encode the α1 and α2 chains of type IV collagen, respectively. These alpha chains interact in the endoplasmic reticulum to form a collagen type IV heterotrimer composed of α1α1α2(IV), which is stabilized mainly by the Gly-X-Y motif alongside other molecular assembly mechanisms^11, 12^. Once secreted, collagen IV proteins self-assemble into a covalently cross-linked lattice style network that connects to the laminin matrix scaffold via bridging molecules, most notably nidogens and heparan sulphate proteoglycans (perlecan)^13^. Type IV collagen is the major structural component of the basement membrane (BM) in blood vessels. Interestingly, in the CNS, there are two functionally distinct BMs, the vascular BM (vBM) and parenchymal (or glial) BM (pBM)^14, 15^. The vBM is synthesised by brain microvascular endothelial cells (BMECs) and vascular mural cells (MCs) and enriched in type IV collagen (α1α1α2(IV)), laminin α4, perlecan (HSPG2), and nidogen^14, 16^. The pBM, in contrast is enriched in laminin α2 and laminin α1 that are secreted by astrocytes, in addition to other ECM proteins^14, 17^.

Astrocytes, the most abundant glial cells in the CNS, are central for regulating neurovascular coupling and BBB function by direct interaction with BMECs in cerebral microvessels and are critical in supporting CNS function by buffering extracellular potassium, clearing and recycling neurotransmitters like glutamate, maintaining metabolic balance, and participating in neuroinflammation. However, the role of astrocyte-secreted pBM components in cSVD development remains unclear and understudied.

The vBM and pBM play important roles in supporting the integrity of the neurovascular unit (NVU) that is composed of multiple neurovascular cell types including BMECs, MCs, astrocytes and neurons. While the vBM provides structural support for BMECs, regulates cell polarity, and anchors tight and adherens junction complexes, which are critical for maintaining blood-brain barrier (BBB) integrity and selectivity^14, 18^, the pBM anchors astrocyte end-feet to microvessels, facilitating bidirectional signalling between the brain parenchyma and circulation. Beyond mechanical stability, the pBM is essential for perivascular homeostasis, influencing immune cell trafficking, ion exchange, and metabolic support to neurons^16, 19, 20^. Patients with pathogenic variants in *COL4A1* or *COL4A2* have a wide spectrum of cSVD associated features including ischemic stroke, spontaneous intracranial hemorrhage (ICH), porencephaly, leukoencephalophy, cerebral microbleeds, and seizures, with radiological finding including white matter hyperintensities (WMHs) and lacunes as part of rare multi-systemic Gould syndrome^21–23^. However, how the mutant collagen leads to these brain pathologies remains unclear.

Using induced pluripotent stem cells (iPSCs) from *COL4A1/COL4A2* cSVD patients, our previous study reported detrimental effects of the mutant MC-secreted vBM in supporting BBB function^24, 25^. Using the same iPSC model reported by Al-Thani et al^25^, this study has focused on the role of astrocytes and astrocyte-secreted pBM components in cSVD pathology. We identified global ECM disorganization in astrocytes that were derived from iPSCs (iPSC-astrocytes) of the *COL4A1^G^*^755^*^R^* cSVD patient. ECM preparations from the mutant astrocytes significantly damaged tight junction (TJs) of iPSC-derived BMECs (iPSC-BMECs), and secretome from the mutant astrocytes reduced its ability to rescue the integrity of TJ in mutant BMECs. Importantly, we identified HTRA1, a serine protease involved in genetic cSVD, to be significantly downregulated in *COL4A1^G^*^755^*^R^* iPSC-astrocytes, and its supplementation rescued the BBB integrity. Our findings provide new insights into the role of astrocytes and the pBM in the regulation of BBB function and driving cSVD pathology, which may inform the development of targeted therapies.

## Results

### Transcriptomics of *COL4A1^G^*^755^*^R^*iPSC-astrocytes highlight ECM defects

The *COL4A1*^G755R^ and isogenic control iPSC lines were previously reported^25^, and their genotype and pluripotency were confirmed in this study (**Fig. S1**). We first differentiated iPSCs into astrocytes using a two-stage protocol involving NPC induction followed by astrocyte differentiation as illustrated in **Figure 1A**. Successful NPC differentiation were confirmed by the rosette morphology and expressions of PAX6 and SOX1 (**Fig. 1B and C**). The subsequent astrocyte differentiation proceeded to Day 60, during which the “astro” shaped astrocytes gradually emerged (**Fig. 1B**). Immunostaining and RT-qPCR demonstrated robust expression of glial fibrillary acidic protein (GFAP) and S100 calcium-binding protein B (S100B), confirming the identity of iPSC-astrocytes (**Fig. 1B and C**).

**Figure 1.**
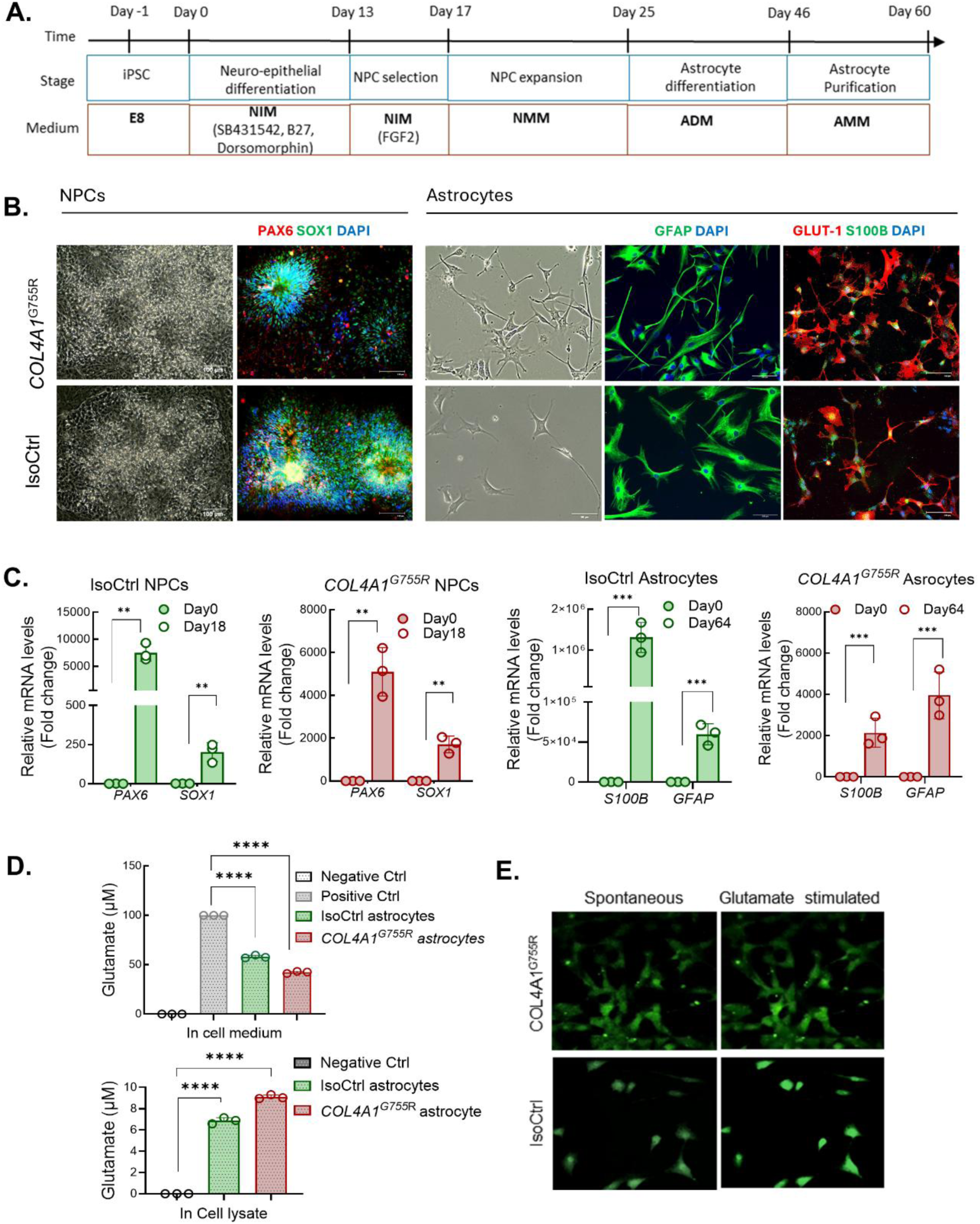
Astrocyte differentiation and characterization. (**A**). Schematic protocol of astrocyte differentiation from iPSCs. (**B**). Phase contrast microscopy and immunofluorescent staining of cells during astrocyte differentiation. NPC markers PAX6 (red) and SOX1 (green) were stained on Day 20, and astrocyte markers GFAP (green), S100B (green) and Glut-1(red) stained on Day 60. Nuclei were stained with DAPI (blue). Scale bars, 100 μm. (**C**). qRT-PCR quantification of NPC and astrocyte specific marker genes *PAX6, SOX1, S100B*, and *GFAP* in isogenic control (IsoCtrl) and *COL4A1*^G755R^ cells on differentiation on Day 0, 18, and 64, respectively. *GAPDH* was used as the endogenous control. (**D**). Glutamate uptake assay. IPSC-astrocytes were incubated with 100 nM glutamate at 37°C for two hours followed by measuring the remaining glutamate in the cell culture medium and cell lysates. Data in (C and D) are mean ± SEM from triplicate reactions of 3 biological replicates, n=3. Unpaired Student’s *t*-test in (C) and one-way ANOVA followed by Tukey’s post hoc test in (D), **p<0.01, ***p<0.001, ****p<0.0001. (**E**). Calcium signals measured by Fluo-4 AM indicator in *COL4A1*^G755R^ and IsoCtrl iPSC-astrocytes. Scale bars, 100 μm. NIM, neural induction medium; NMM, neural maintenance medium; ADM, astrocyte differentiation medium; AMM, astrocyte maturation medium.

To evaluate the functional maturity of iPSC-astrocytes, we performed glutamate uptake assay (**Fig. 1D**). Cells were exposed to a defined amount of glutamate for two hours followed by colorimetric quantification of glutamate in both cell lysates and medium, where increased cellular glutamate and concomitant depletion from the medium indicated active glutamate uptake. Astrocytes derived from both the mutant and isogenic control iPSCs had the ability of up-taking glutamate from the medium (**Fig. 1D**). Notably, *COL4A1*^G755R^ astrocytes had significantly enhanced glutamate uptake compared to the isogenic control (**Fig. 1D**), suggesting a potential metabolic stress or a reactive phenotype. Calcium imaging revealed spontaneous baseline calcium oscillation in the cells and responses to glutamate stimulation (**Fig. 1E** and **S2**). The assessments on cell morphology, marker gene expression and the functional assay validated the iPSC-astrocytes for modelling COL4A1-related cellular phenotypes.

To identify molecular mechanisms of *COL4A1*-related cSVD, a global transcriptomic approach was conducted using RNA sequencing (RNAseq). PCA plot revealed a clear separation between samples of *COL4A1*^G755R^-astrocytes and isogenic controls on PC1 that explains 67% of the total variance, indicating a distinct transcriptomic signature of the mutant astrocytes **(Fig. 2A).** Data analysis identified 1,284 significantly DEGs between the two groups, of which 833 were upregulated and 451 downregulated **(Fig. 2B)**. GO analysis revealed significant extracellular matrix-related alterations across all three major GO categories (**Fig. 2C**). The most significantly enriched GO terms were “extracellular matrix organization” and “extracellular structure organization” under the biological process (BP) category, underscoring a central defect in ECM pathways in mutant astrocytes. We then conducted GSEA analysis to uncover the coordinated changes of genes involved in ECM function. Results show that gene sets involved in “collagen fibril organization” and “regulation of extracellular matrix organization” were negatively enriched (**Fig. 2D**).

**Figure 2.**
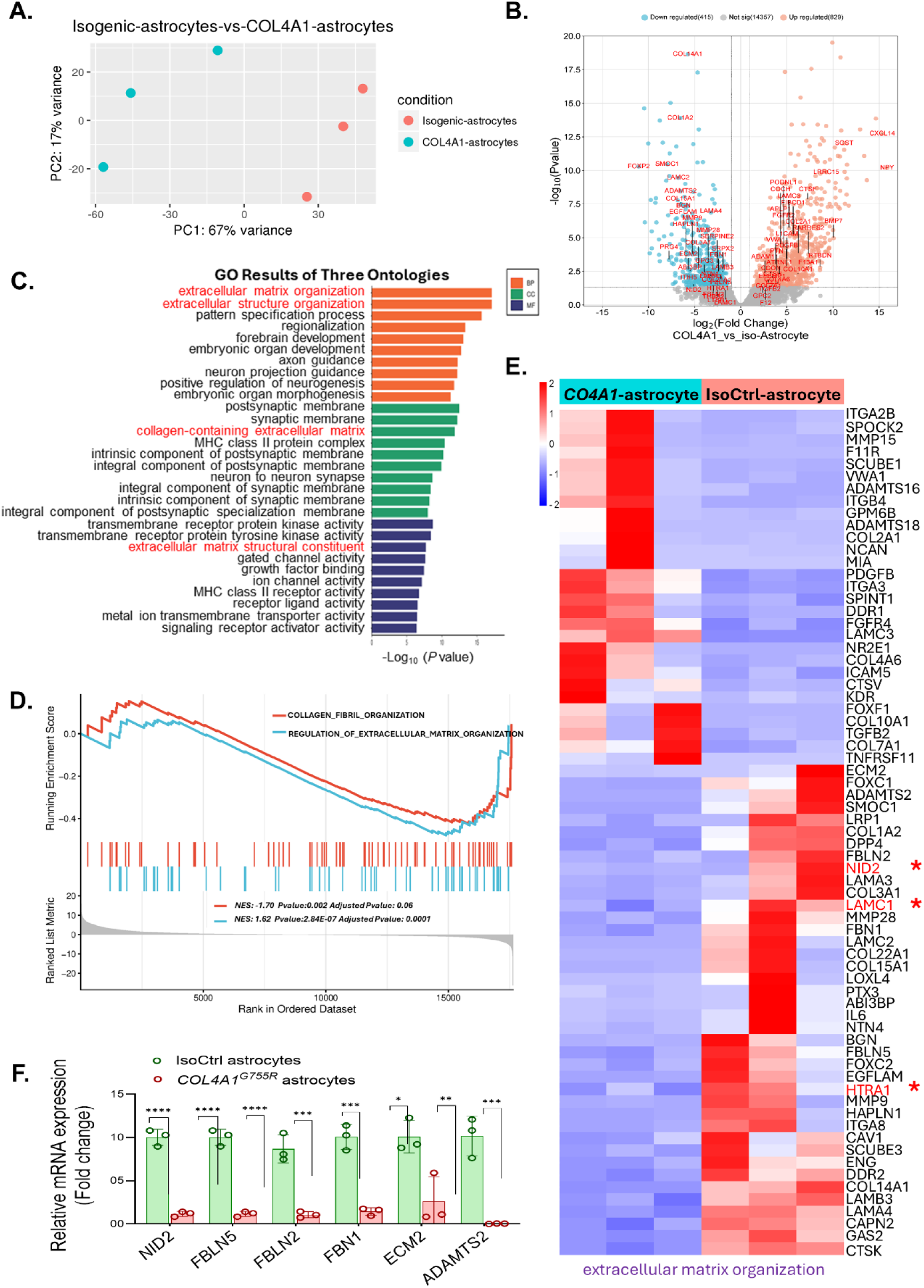
Transcriptome Profiling of iPSC-astrocytes. Astrocytes derived from *COL4A1*^G755R^ and isogenic control iPSCs from 3 independent iPSC differentiations were subjected to RNA sequencing (RNA-seq) followed by data analysis. (**A**). Principal component analysis (PCA). (**B**). Volcano plot displaying differentially expressed genes (DEGs) in *COL4A1*^G755R^ astrocytes compared to the isogenic control with an adjusted p-value cutoff p<0.05. Log2 fold change greater than 1 are indicated by red dots, representing significantly up-regulated DEGs; and log2 fold change less than -1 are indicated by blue dots, representing significantly down-regulated DEGs. (**C**). Gene ontology (GO) analysis of DEGs demonstrates enriched GO terms of biological process (BP), cellular components (CC), and molecular functions (MF). (**D**). GSEA plot of DEGs revealing enriched gene sets relating to the indicated GO terms. (**E**). Heatmap showing DEGs of *COL4A1*^G755R^ and isogenic control astrocytes under the GO term “extracellular matrix organization”. (**F**). RT-qPCR conformation of RNAseq results on basement membrane related genes. Data are means ± SEM from triplicate reactions of 3 biological replicates, n=3. Unpaired Student’s *t*-test, **p<0.01, **p<0.01, ***p<0.001, ****p<0.0001.

### Defects of pBM in *COL4A1*^G755R^ iPSC-astrocytes and associated HTRA1 deficiency and TGF-β signalling dysregulation

To further dissect ECM alterations in the *COL4A1*^G755R^ astrocytes, we analysed DEGs under the GO term “extracellular matrix organization” as shown on the heatmap (**Fig. 2E**), and these DEGs were also annotated on the volcano plot (**Fig. 2B**). The DEG list contains key genes encoding pBM proteins secreted by astrocytes including *LAMC1, NID2*, *FBLN2*, *FBLN5*, *FBN1,* and ECM remodeling genes such as *MMP9* and *HTRA1*, most of which were downregulated. We then verified the RNAseq results for some of these genes using RT-qPCR (**Fig. 2F**). Western blotting (WB) confirmed that Nidogen 2 (encoded by *NID2*), the pBM-specific nidogen type, was significantly reduced in *COL4A1*^G755R^ astrocytes (**Fig. S3A**), and immunofluorescent staining of ECM secreted from the *COL4A1*^G755R^ astrocytes revealed sparse nidogen signal, comparing to the isogenic control (**Fig. S3B**). *COL4A1* expression was not significantly altered (**Fig. S3C**), since the *COL4A1*^G755R^ variant is predicted to affect posttranscriptional assembly of the collagen triple-helix; however, its secretion to extracellular space tends to be compromised (**Fig. S3B**), which is in line with previous findings^25^.

Among the differentially expressed pBM genes, *HTRA1* attracted our specific interest as it is a causative gene for monogenic cSVD, homozygous mutations resulting in CARASIL and heterozygous mutations causing a milder clinical phenotype in CADASIL-2^26, 27^. HTRA1 is a serine protease that plays a key role in maintaining BM homeostasis by degrading misfolded ECM proteins and regulating TGF-β activity^28^. Using RT-qPCR and WB we confirmed a significant down regulation of HTRA1 in *COL4A1*^G755R^ astrocytes (**Fig. 3A and B**). Immunofluorescence staining of both iPSC-astrocytes and ECM preparations showed less abundant HTRA1 around the *COL4A1*^G755R^ astrocytes and in the ECM (**Fig. 3C**). Concomitantly, a marked increase of ECM-associated TGF-β1 deposition was observed in the ECM preparation from *COL4A1*^G755R^ astrocytes compared to the isogenic control (**Fig. 3D**), suggesting increased extracellular availability of TGFβ1 and potential activation of TGF-β signaling. This was confirmed by phosphorylated Smad2/3 (P-Smad2/3) staining which demonstrated a distinct shift of P-Smad2/3 from cytoplasmic to predominant nuclear localisation in *COL4A1*^G755R^ astrocytes (**Fig. 3E and F**), a hallmark of canonical TGF-β signaling activation. RNAseq analysis also highlighted increased expression of *TGFB2* (**Fig. 2E**), and RT-qPCR confirmed the elevation of both *TGFB1* and *TGFB2*, and decreased *LTBP1* (**Fig. 3G**).

**Figure 3.**
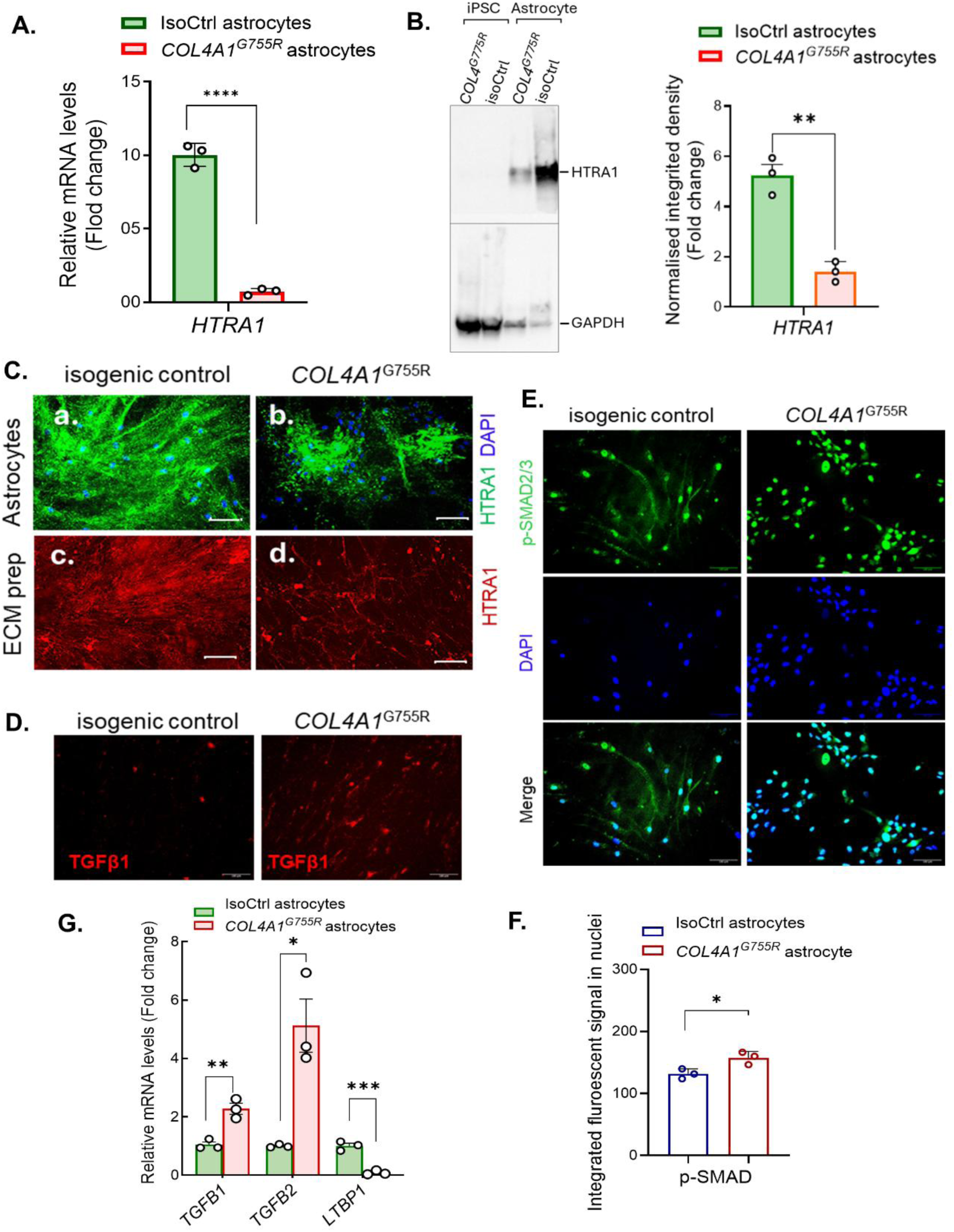
Changes of HTRA1 in *COL4A1^G^*^755^*^R^*iPSC-astrocytes and dysregulation of TGF-β signalling. Astrocytes were differentiated from *COL4A1*^G755R^ and isogenic control (IsoCtrl) iPSCs and subjected to RT-qPCR and immunofluorescent staining. (**A**). *HTRA1* expression determined by RT-qPCR. (**B**). HTRA1 protein levels determined by western blotting (WB). WB of iPSC samples of *COL4A1*^G755R^ and IsoCtrl were also included as controls. Right panel shows quantification of WB images. (**C**). Immunofluorescent staining of HTRA1 in iPSC-astrocytes (a and b) and ECM preparations from astrocytes (c and d). (**D**). Immunofluorescent staining of TGF-β in ECM preparations of iPSC-astrocytes. (**E**). Immunofluorescent staining of p-SMAD2/3 in iPSC-astrocytes. (**F**) Quantification of nuclear signals of the p-SMAD2/3 staining in (E). (**G**). *TGFB1, TGFB2, and LTBP1* expression in iPSC-astrocytes were determined by RT-qPCR. Data in (A, B, F and G) are means ± SEM from triplicate reactions of 3 biological replicates, n=3. Unpaired Student’s *t*-test, **p<0.01, **p<0.01, ***p<0.001, ****p<0.0001. Scale bar in (C-E), 100 μm.

### ECM from *COL4A1*^G755R^ iPSC-astrocytes damages tight-junction integrity of iPSC-BMECs

As the pBM is a key regulator of BMEC function and BBB integrity, we evaluated the direct effect of the *COL4A1*^G755R^ astrocyte-secreted ECM on BMECs. We differentiated iPSCs into BMECs (iPSC-BMECs) using a protocol modified from our previous publication^29^ and the publication by Pediaditakis et al ^30^ (**Fig. S4A-C**). We first confirmed previous findings where TJs in *COL4A1*^G755R^ BMECs were significantly damaged as stained for claudin 5, occluding and ZO-1, and TEER was also significantly reduced compared to the isogenic control (**Fig. S4D and E**)^25^. We then showed that isogenic control BMECs grown on the ECM from *COL4A1*^G755R^ mutant astrocytes developed significantly disrupted TJ architecture characterised by irregular, discontinuous, and punctate staining patterns, compared to control BMECs growing on ECM preparations from the control astrocytes (**Fig. 4A and B**). We then grew the *COL4A1*^G755R^ mutant BMECs on ECM preparations of the isogenic control or mutant iPSC-astrocytes and found that the control astrocyte ECM significantly restored the TJ integrity of the mutant BMECs (**Fig. 4C and D**), suggesting the importance of a healthy pBM in supporting BBB function.

**Figure 4.**
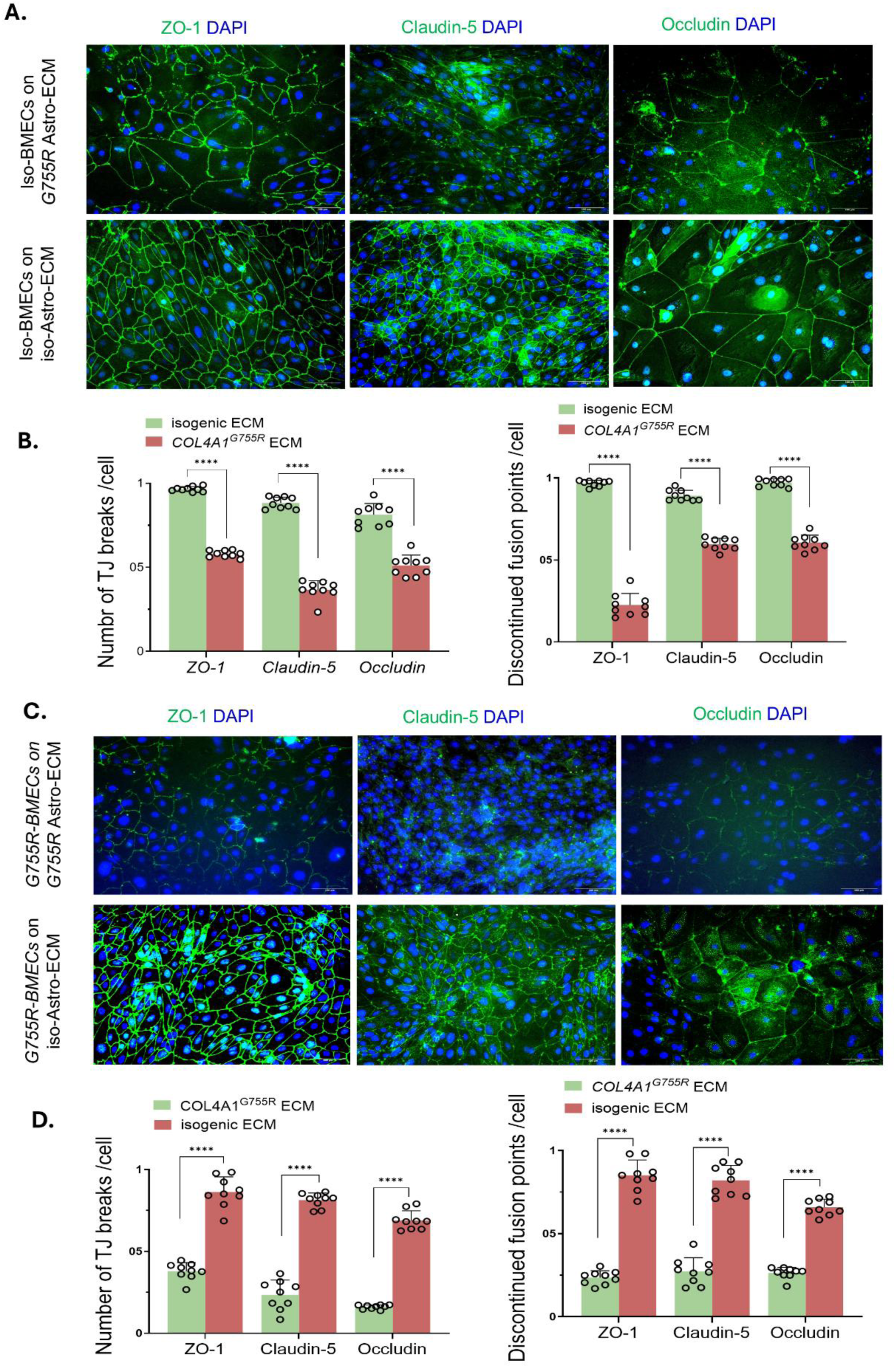
ECM secreted from *COL4A1^G^*^755^*^R^* iPSC-astrocytes damages tight junctions formed by iPSC-BMECs. IPSC-BMECs were seeded onto extracellular matrix (ECM) synthesised from iPSC-astrocytes for 4 days, followed by immunofluorescent staining of tight junction (TJ) markers ZO-1, Claudin-5, and Occludin. (**A**). Isogenic BMECs were grown on ECM secreted from isogenic and *COL4A1^G^*^755^*^R^*astrocytes. (**B**). Qualifications of (A) on the integrity of TJs. Figure shows the number of TJ breaks per cell (left panel) and discontinued fusion points per cell (right panel). (**C**). *COL4A1^G^*^755^*^R^* BMECs were grown on ECM secreted from isogenic and *COL4A1^G^*^755^*^R^*astrocytes. (**D**). Qualifications of (B) on the integrity of TJs. Figure shows the number of TJ breaks per cell (left panel) and discontinued fusion points per cell (right panel). Data in (B and D) are presented as means ± SEM from 9 images, with 3 randomly selected images taken from each of 3 independent experiments, n=9. Unpaired Student’s *t*-test, **** p<0.0001.

### iPSC-astrocyte secretome enhances BMEC barrier function with reduced efficacy from *COL4A1*^G755R^ iPSC-astrocytes

To better understand the contribution of astrocyte-derived soluble factors, in addition to ECM, to the disease mechanism in *COL4A1*-associated cSVD, we investigated the effects of the astrocyte secretome on BBB integrity. We cultured isogenic control iPSC-BMECs in conditioned medium from either the mutant *COL4A1*^G755R^ iPSC-astrocytes or the isogenic control iPSC-astrocytes in a Trans-well insert for four days. Results showed that while the TJ continuity of the control BMECs were not further enhanced by either the control or mutant astrocyte secretome (**Fig. 5A and B**), both the control and mutant astrocyte secretomes induced significantly increased protein levels of ZO-1, occludin and claudin-5 in control BMECs, but this effect was significantly weaker by the mutant astrocyte secretome (**Fig. 5C**). These results suggest that the secretome of astrocytes carrying the *COL4A1*^G755R^ mutation did not completely lose their ability to support BMEC TJ integrity, but their supportive capacity is compromised. Notably, the mRNA expression of *ZO-1, CLND5* and *OCDN* in control BMECs were unchanged when cultured in conditioned medium from either the control or mutant astrocytes (**Fig. 5E**), suggesting a posttranslational mechanism.

**Figure 5.**
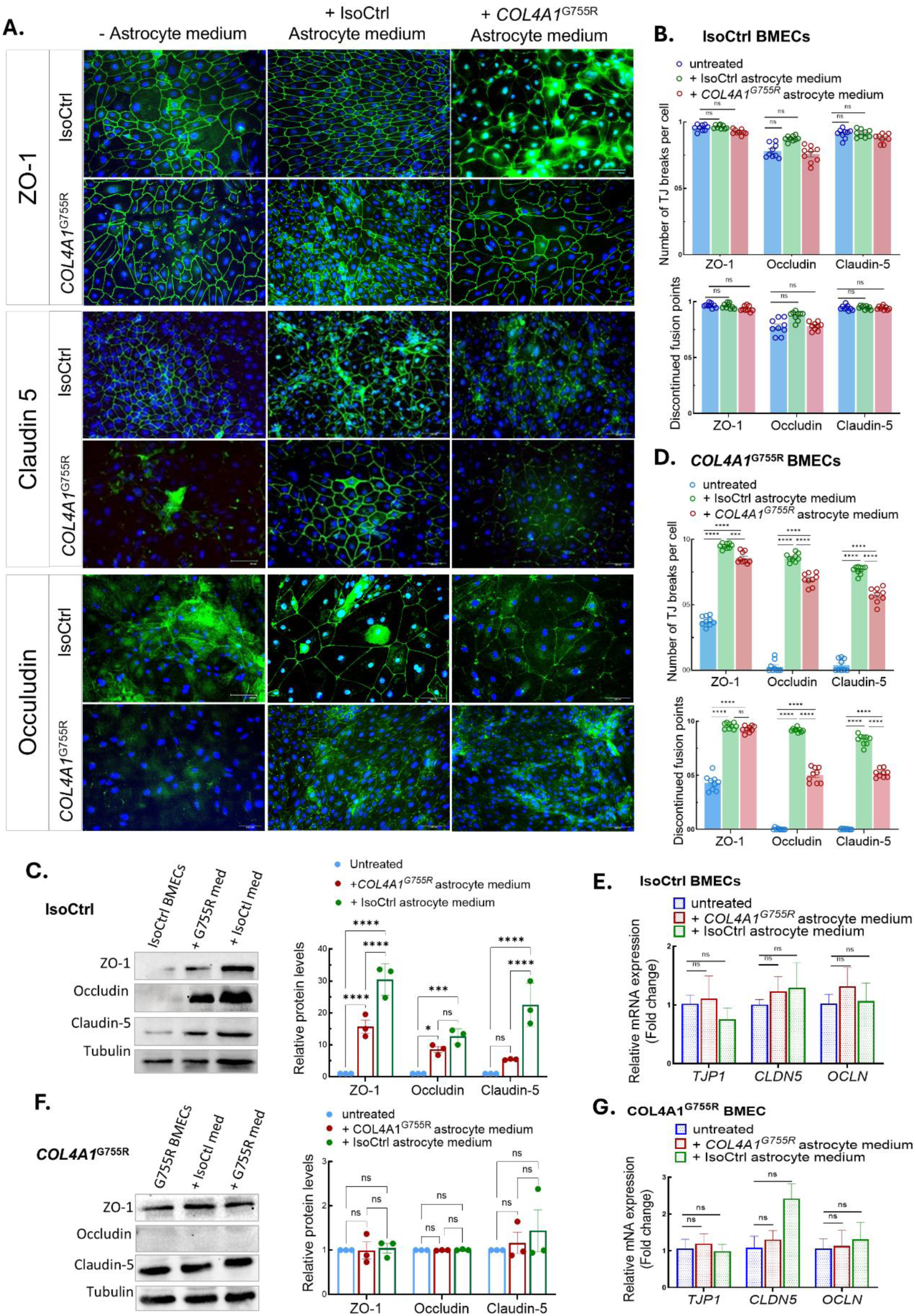
Effect of iPSC-astrocyte secretome on tight junction integrity in iPSC-BMECs. BMECs and astrocytes were differentiated from iPSCs of *COL4A1*^G755R^ and its isogenic control (IsoCtrl). Total RNA and cells were collected from *COL4A1^G^*^755^*^R^*iPSC-BMECs after 4 days of treatment with astrocyte-conditioned medium (ACM) from patient or control iPSC-astrocytes, mixed at a ratio of ACM: hECSR2 = 7:3. Protein of *COL4A1^G^*^755^*^R^*iPSC-BMECs were collected from a Trans-well® co-culture system after co-cultured with iPSC-astrocytes for 4 days. (**A**). Immunofluorescent staining of tight junction (TJ) markers ZO-1, Claudin-5, and Occludin in *COL4A1*^G755R^ or IsoCtrl iPSC-BMECs cultured without astrocyte medium (left column), with IsoCtrl astrocyte medium (middle column), or with *COL4A1*^G755R^ astrocyte medium (right column), respectively. Scale bar, 100 μm. (**B**). Qualifications of (A) on the TJ integrity in IsoCtrl BMECs exposed to either IsoCtrl or *COL4A1*^G755R^ astrocyte conditioned medium. Top figure, the number of TJ breaks per cell; bottom figure, discontinued fusion points per cell. (**C**) WB of TJ proteins ZO-1, Claudin-5, and Occludin in IsoCtrl iPSC-BMECs co-culture with either IsoCtrl or *COL4A1*^G755R^ astrocyte. Quantifications of band densities are shown on the right panel. (**D**). Qualifications of (A) on the TJ integrity in *COL4A1*^G755R^ BMECs exposed to either IsoCtrl or *COL4A1*^G755R^ astrocyte medium. Top figure, the number of TJ breaks per cell; bottom figure, discontinued fusion points per cell. (**E**). RT-qPCR results of IsoCtrl BMECs exposed to either IsoCtrl or *COL4A1*^G755R^ astrocyte conditioned medium. (**F**). WB of TJ proteins ZO-1, Claudin-5, and Occludin in *COL4A1*^G755R^ iPSC-BMECs co-culture with either IsoCtrl or *COL4A1*^G755R^ astrocyte. Quantifications of band densities are shown on the right panel. (**G**). RT-qPCR results of *COL4A1*^G755R^ BMECs exposed to either IsoCtrl or *COL4A1*^G755R^ astrocyte conditioned medium. Data are means ± SEM from triplicate reactions of 3 biological replicates, n=3. Two-way ANOVA followed by Šídák’s post hoc test, **p<0.01, **p<0.01, ***p<0.001, ****p<0.0001.

We next examined the effects of secretomes from the isogenic control and mutant iPSC-astrocyte on the TJ integrity of the *COL4A1*^G755R^ mutant BMECs using the same Trans-well co-culture setting. Both secretomes could significantly restore the disorganised TJs of the mutant BMECs, but again, the extent of the restoration by the mutant astrocyte secretome was significantly compromised compared to that of the control (**Fig. 5A and D**). However, neither the healthy or mutant astrocyte secretomes increased TJ protein levels in *COL4A1*^G755R^ mutant BMECs, suggesting an intrinsic or autonomous defect on the regulation of TJ proteins in mutant BMECs (**Fig. 5F**). This finding, taking the TJ-rescue capability by the mutant astrocyte secretome (**Fig. 5A and D**) into account, indicates that the observed improvement in TJ organisation is likely mediated by redistribution or reorganisation of existing TJ proteins rather than changes in their overall levels. Accordingly, the mRNA expression of *ZO-1, CLND5* and *OCDN* remained unchanged in BMECs when cultured in conditioned medium from either the mutant or control astrocytes (**Fig. 5G**).

#### COL4A1^G755R^ iPSC-astrocyte secretome exacerbates the ECM dysregulation in COL4A1^G755R^ BMECs

To model the disease-relevant microenvironment for *COL4A1*-related cSVD, we performed bulk RNAseq on *COL4A1^G^*^755^*^R^*iPSC-BMECs cultured in conditioned medium from *COL4A1^G^*^755^*^R^* iPSC-astrocytes, alongside isogenic control iPSC-BMECs exposed to isogenic control astrocyte-conditioned medium. This comparison identified 656 DEGs (395 upregulated and 261 downregulated; **Fig. 6A**). In contrast, monocultured iPSC-BMECs, i.e., BMECs grown without astrocyte secretomes, showed a substantially larger transcriptomic divergence, especially the downregulated DEGs (300 upregulated and 663 downregulated; **Fig. 6B**). These findings indicate that both control and mutant astrocyte secretomes reduce global transcriptomic variability between the mutant and control iPSC-BMECs, consistent with our early observations where astrocyte secretomes exert a stabilising effect on BMECs (**Fig. 5A and C**).

**Figure 6.**
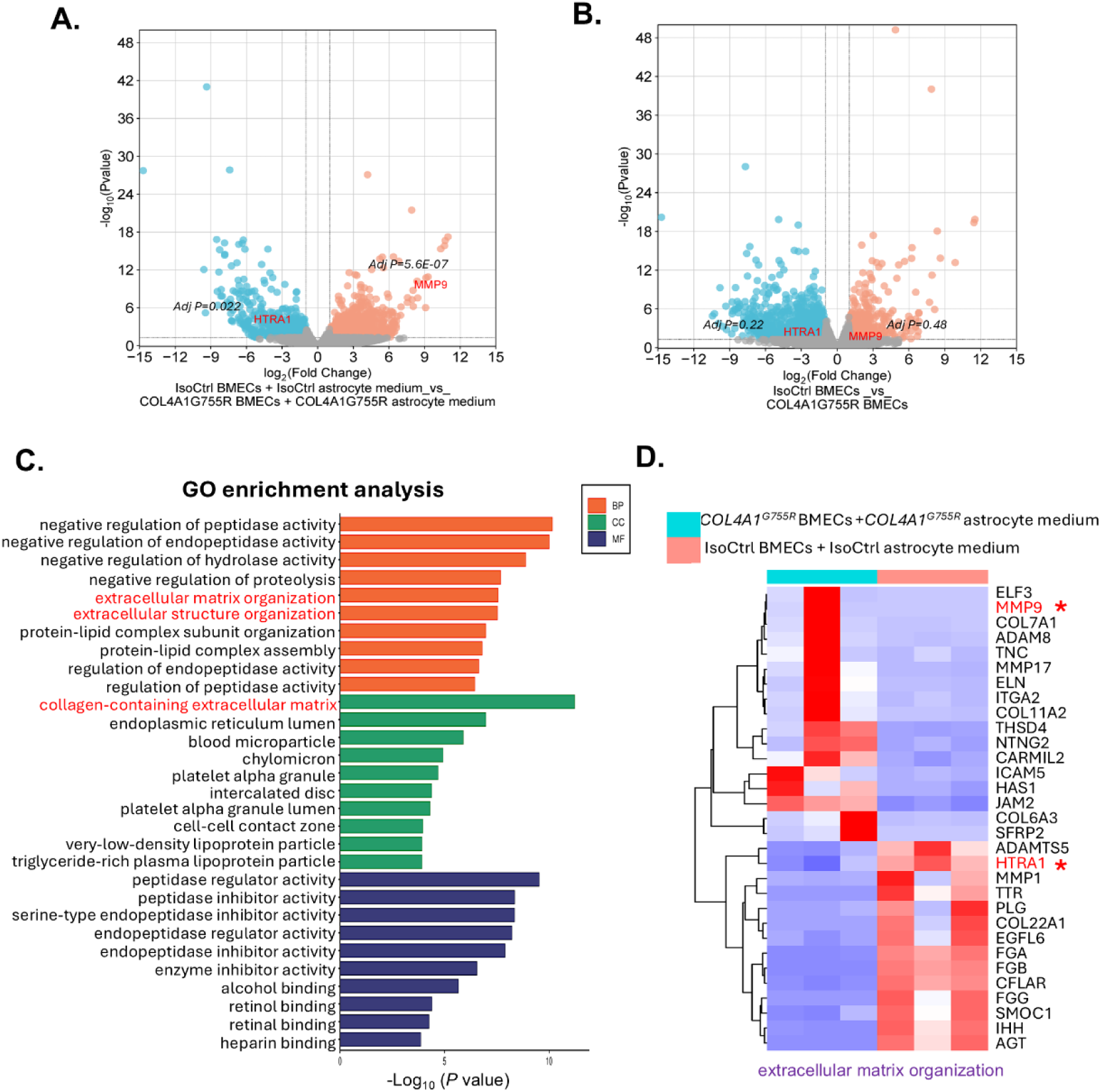
Transcriptomics of iPSC-BMECs exposed to secretome of iPSC-astrocytes. BMECs and astrocytes were differentiated from iPSCs of *COL4A1*^G755R^ and its isogenic control (IsoCtrl) from 3 independent experiments. The iPSC-derived *COL4A1*^G755R^ and IsoCtrl BMECs were then cultured in their corresponding iPSC-astrocyte conditioned medium mixed with hECSR2 medium in a 7:3 ratio for 4 days, followed by RNAseq. RNAseq were also conducted on *COL4A1*^G755R^ and IsoCtrl iPSC-BMECs that were not exposed to astrocyte conditioned medium for comparison. (**A-B**). Volcano plots showing differentially expressed genes (DEGs) between *COL4A1*^G755R^ BMECs exposed to *COL4A1*^G755R^ astrocyte conditioned medium and IsoCtrol BMECs exposed to IsoCtrl astrocyte conditioned medium (**A**), and between *COL4A1*^G755R^ BMECs and IsoCtrol BMECs (**B**). (**C**). Gene ontology (GO) analysis of DEGs between *COL4A1*^G755R^ BMECs exposed to *COL4A1*^G755R^ astrocyte conditioned medium and IsoCtrol BMECs exposed to IsoCtrl astrocyte conditioned medium. (**D**). Heatmap showing DEGs under the GO term “extracellular matrix organisation”. Results comparing DEGs between *COL4A1*^G755R^ BMECs exposed to *COL4A1*^G755R^ astrocyte conditioned medium and IsoCtrol BMECs exposed to IsoCtrl astrocyte conditioned medium from 3 independent experiments.

GO enrichment analysis of mutant and control iPSC-BMECs exposed to their corresponding mutant and control astrocyte-conditioned medium revealed significant changes in ECM-related terms, including “extracellular matrix organisation”, “extracellular structure organisation”, and “collagen-containing extracellular matrix” (**Fig. 6C**). These ECM-associated signatures closely mirrored those observed in *COL4A1*^G755R^ astrocytes (**Fig. 2C**), suggesting coordinated matrix dysregulation across cell types. Notably, these ECM-related GO terms were not significantly enriched in monocultured BMECs (**Fig. S5A**), highlighting the importance of astrocyte-derived cues in unmasking disease-relevant transcriptional changes.

Among the ECM-associated DEGs, *HTRA1* emerged as a significantly downregulated transcript in mutant BMECs exposed to mutant astrocyte medium (**Fig. 6A and D**). RT-qPCR and western blotting confirmed this reduction (**Fig. S6A-C**). Although *HTRA1* showed only a non-significant downward trend in monocultured BMECs (adjusted *P* = 0.22; **Fig. 6B**), independent RT-qPCR experiments detected a significant decrease (**Fig. S6B**). Importantly, mutant astrocyte-conditioned medium amplified *HTRA1* downregulation from ∼29-fold to ∼48-fold, indicating that mutant astrocyte secretions exacerbate HTRA1 suppression in BMECs.

Similarly, *MMP9* was significantly upregulated in BMECs exposed to mutant astrocyte medium (**Fig. 6A and S6D**), suggesting enhanced ECM degradation and potential BBB vulnerability. MMP9 was not significantly altered in monocultured BMEC RNA-seq data (**Fig. 6B**), yet RT-qPCR confirmed increased expression (**Fig. S6E**). Mutant astrocyte medium further intensified this change, increasing *MMP9* expression from ∼72-fold to ∼327-fold.

Together, these findings demonstrate that *COL4A1^G^*^755^*^R^* astrocytes actively drive ECM pathology in BMECs and synergise with cell-autonomous *COL4A1* defects to amplify vascular dysfunction. The results also underscore the necessity of multicellular models for accurately capturing cSVD mechanisms.

### HTRA1 treatment rescue impaired tight junctions of *COL4A1*^G755R^ iPSC-BMECs and improved astrocyte function

Given the observed downregulation of HTRA1 in *COL4A1*^G755R^ iPSC-astrocytes and BMECs and its established role in ECM remodelling and signalling regulation, as well as its involvement in other genetic cSVD, we hypothesised that correction of HTRA1 deficiency could rescue the disruption of TJ integrity, a hallmark feature of BBB dysfunction in *COL4A1*-related cSVD. To test this hypothesis, recombinant human HTRA1 (20 ng/ml) was administered to *COL4A1*^G755R^ iPSC-BMECs for 48 hours. Immunofluorescence analysis revealed that HTRA1 treatment significantly rescued the TJ architecture (**Fig. 7**). The untreated *COL4A1*^G755R^ BMECs displayed fragmented and discontinuous TJ patterns of TJ proteins including ZO-1, occludin, and claudin-5, and often interrupted at fusion points. In contrast, HTRA1 treated cells exhibited a more uniform and linear staining pattern (**Fig. 7A**).

**Figure 7.**
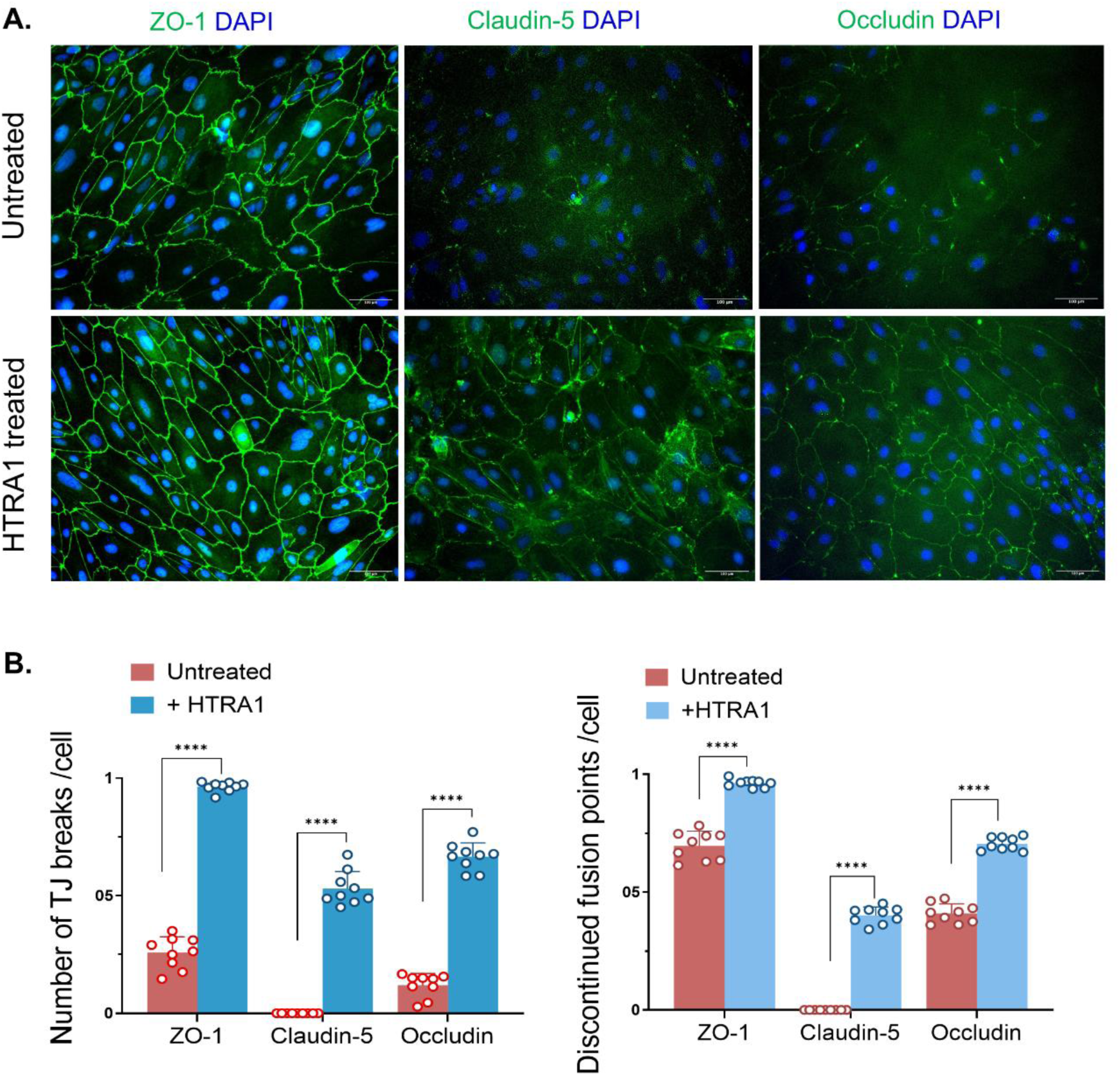
HTRA1 treatment rescue impaired tight junctions of *COL4A1*^G755R^ iPSC-BMECs. BMECs derived from *COL4A1*^G755R^ iPSCs were treated with or without human HTRA1 recombinant protein (20 ng/mL) for 4 days, with medium refreshed every 48 hours. The cells were then subjected to immunofluorescent staining for tight junction (TJ) markers ZO-1, Claudin-5 and Occludin. (**A**). Immunofluorescent images of TJs. Scale bar, 100 μm. (**B**). Quantification of TJ continuity in (A) which are presented as the number of TJ breaks per cell (left panel) and discontinued fusion points per cell (right panel). Data are means ± SEM derived from 9 images, with 3 randomly selected images taken from each of 3 independent experiments, n=9. Unpaired Student’s *t*-test, ****p<0.0001.

To further investigate the mechanism underlying the protective role of BBB by HTRA1, we investigated the involvement of TGFβ signalling as hypothesised earlier. RT-qPCR analysis showed that HTRA1 supplementation significantly suppressed the elevated mRNA expression of *TGFB1*, *TGFB2*, and *TGFBR2* in *COL4A1*^G755R^ iPSC-BMECs (**Fig. 8A)**. We then tested if TGF-β receptor inhibitor could replicate the rescue effect of HTRA1 by applying SB431542 (10 μM) to the *COL4A1^G^*^755^*^R^* iPSC-BMEC culture. Immunofluorescence analysis demonstrated that SB431542 treatment significantly improved TJ organization, as evidenced by the improved continuity of ZO-1 and occludin staining at cell boundaries and fusion points (**Fig. 8B-D**).

**Figure 8.**
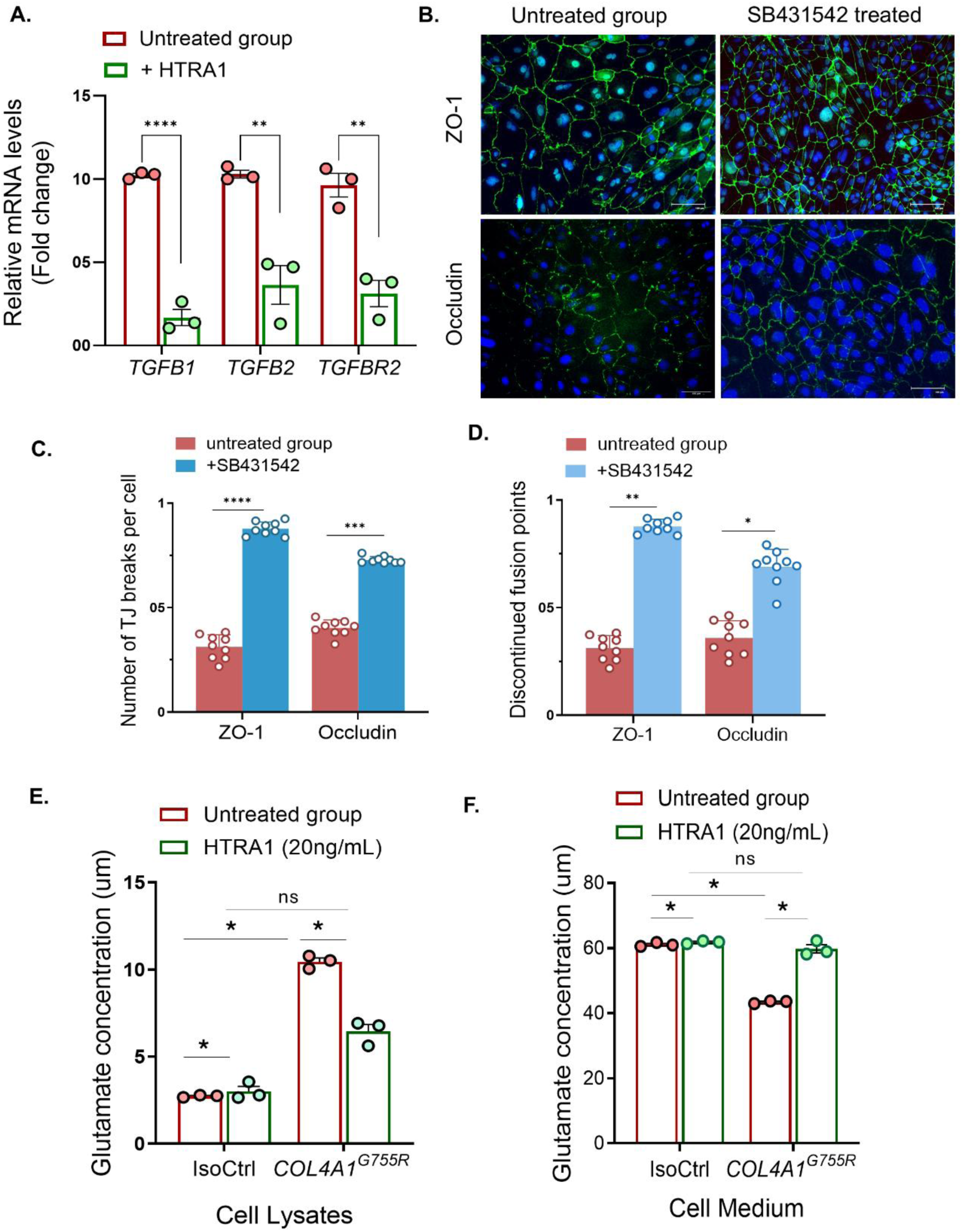
Involvement of TGF-β signalling in HTRA1 rescue of impaired tight junctions in *COL4A1*^G755R^ iPSC-BMECs. (**A**). BMECs derived from *COL4A1*^G755R^ iPSCs were treated with or without human HTRA1 recombinant protein (20 ng/mL) for 4 days, with medium refreshed every 48 hours. RT-qPCR showing reduced expression of *TGFB1, TGFB2* and *TGFR2* after HTRA1 treatment. Data are means ± SEM, n=3. Unpaired Student’s *t*-test, **p<0.01, ***p<0.001, ****p<0.0001. (**B**). BMECs derived from *COL4A1*^G755R^ iPSCs were treated with or without TGF-β inhibitor SB431542 (10 μM) for 4 days, followed by immunofluorescent staining of tight junction (TJ) proteins ZO-1 and Occludin. Scale bar, 100 μm. (**C-D**). Quantification of TJ continuity in (A) which are presented as the number of TJ breaks per cell (C) and discontinued fusion points per cell (D). Data are means ± SEM derived from 9 images, with 3 randomly selected images taken from each of 3 independent experiments, n=9. Unpaired Student’s *t*-test, *p<0.05, **p<0.01, ***p<0.001, ****p<0.0001. (**D**). Astrocytes derived from *COL4A1*^G755R^ and isogenic control (IsoCtrl) iPSCs were treated with or without human HTRA1 protein (20 ng/mL), respectively, for 4 days, with medium refreshed every 48 hours. Glutamate uptake was then measured. Data are means ± SEM from triplicate reactions of 3 biological experiments, n=3. Unpaired Student’s *t*-test, *p<0.05.

Next, we showed that supplementation of recombinant human HTRA1 protein to astrocyte culture almost fully reversed the abnormally enhanced glutamate uptake by the *COL4A1*^G755R^ iPSC-astrocytes (**Fig. 8E and F**). HTRA1 treatment did not significantly change glutamate uptake in the isogenic control iPSC-astrocytes (**Fig. 8F**). In contrast, inhibition of TGF-β signalling by SB431542 resulted in a significant enhancement of glutamate uptake in both *COL4A1*^G755R^ and the isogenic control iPSC-astrocytes (**Fig. S7**), suggesting an alternative mechanism to TGF-β signalling underlies the damaged astrocyte function.

Together, these findings suggest that reduced HTRA1 expression in *COL4A1*^G755R^ astrocytes and BMECs contributes to BBB dysfunction at least in part via upregulation of TGF-β signalling. Restoration of HTRA1 activity or pharmacological inhibition of the TGF-β pathway may therefore represent promising therapeutic strategies for *COL4A1*-related cSVD.

## Discussion

Using an iPSC model derived from a *COL4A1* cSVD patient, this study examined how the *COL4A1*^G755R^ variant disrupts astrocyte function and compromises BBB integrity. Transcriptomic profiling revealed ECM dysregulation in *COL4A1*^G755R^ iPSC derived astrocytes compared with isogenic controls, underscoring the role of the pBM in COL4A1 cSVD pathology. Notably, HTRA1 was downregulated in both *COL4A1*^G755R^ iPSC-astrocytes and BMECs, and supplementation with human recombinant HTRA1 restored BBB integrity and astrocyte function, highlighting a potential therapeutic target for this disease group.

Linking HTRA1 to *COL4A1* cSVD highlights an important disease mechanism. HTRA1 is a secreted serine protease that directly cleaves ECM proteins including fibronectin, aggrecan, collagens, and perlecan^28^. Under physiological conditions, HTRA1 contributes to ECM remodelling during development, degrades aged and damaged ECM components, and helps release ECM-embedded bioactive molecules, acting as a master regulator of ECM homeostasis^28^. HTRA1 has established roles in other genetic cSVDs: biallelic variants cause CARASIL characterized by early-onset ischemic strokes and progressive cognitive deterioration with spondylosis and alopecia^31^; while heterozygous variants cause CADASIL-2 (cerebral autosomal dominant arteriopathy with subcortical infarcts and leukoencephalopathy-2)^32^, the second most common hereditary cSVD. Interestingly, rare pathogenic *HTRA1* variants are also found in the general population, including the UK Biobank, which increase the risk of stroke and dementia^33^, and common variants (SNPs) in *HRTA1* and *COL4A1* associate with white matter hyperintensity (WMH) burden^34–36^. Most cSVD-associated *HTRA1* variants are loss-of-function. Supporting this, a recent large proteomics study of >40,000 individuals with MRI markers of cSVD found that lower plasma HTRA1 levels associated with dementia and hippocampal perivascular spaces^37^, and loss-of-protease function of *HTRA1* variants associated with ischemic stroke and coronary artery disease^38^. Reduced HTRA1 has also been observed in brain vessels of CADASIL^26^, the most prevalent genetic cSVD caused by *NOTCH3* variants, and CAA, a common sporadic cSVD and age-related dementia, where HTRA1 becomes sequestered into aggregated NOTCH3 extracellular domain (N3ECD) and in Aβ-containing microvascular deposits^27, 39^, suggesting additional mechanisms leading to HTRA1 loss-of-function across cSVDs.

Researchers have long been seeking a convergent mechanism to explain diverse forms of cSVD and that might reveal shared therapeutic targets. Although the matrisome has been proposed to be a promising candidate^10^, a more specific gene or pathway remains elusive. Based on literature findings described above, HTRA1 seems to be a specific target, however, among the most common genetic cSVDs caused by gene variants such as *NOTCH3*, *HTRA1* and *COL4A1/A2*, there has not been direct evidence demonstrating the contribution of HTRA1 to pathologies specific to COL4A1/A2 disease. In this study, we provide experimental evidence where HTRA1 was significantly downregulated in astrocytes of a patient-specific iPSC model of *COL4A1* cSVD, suggesting HTRA1 inactivation is likely a promising convergent mechanism contributing to the development of the common types of genetic, and possibly sporadic, cSVD. Most importantly, treatment of the iPSC COL4A1 cSVD model with human recombinant HTRA1 significantly rescued the impaired TJs of *COL4A1*^G755R^ iPSC-BMECs, reversed the BBB integrity, and improved astrocyte function, representing the first HTRA1-targeted intervention in cSVD models. However, how the *COL4A1* mutation drives HTRA1 downregulation in astrocytes remains unclear. HtrA1 is a specific astrocyte marker in adult mouse forebrain and plays an important role in astrocyte development^40^. HtrA1 expression is regulated by BMP4 signalling in postnatal mouse astrocytes and responses to brain injury^40^. Defining the mechanisms underlying reduced HTRA1 in *COL4A1* cSVD will be essential for our future work.

An important function of HTRA1 is regulating the TGF-β signalling activity. It has been established that HTRA1 negatively regulates TGF-β signalling through proteolytic degradation of the latency-associated peptide (LAP)^41–43^. Consistent with this role, *COL4A1*-mutant astrocytes in our model showed reduced HTRA1 alongside increased TGF-β signalling components, including elevated ECM-associated TGF-β1 and nuclear P-Smad2/3. These findings align with reports of heightened TGF-β activity, TGF-β1 protein accumulation, and increased Smad2/3 expression in human samples and a mouse model of CARASIL^42–44^. In contrast, another CARASIL mouse model reported reduced TGF-β signalling due to loss of HtrA1-mediated cleavage of latent TGF-β1 binding protein-1 (LTBP-1)^45^, suggesting that TGF-β regulation may be context- or cell-type-dependent. Supporting our observations, a *Col4a1* mutant mouse model of Gould syndrome also showed elevated TGF-β signalling, and its suppression prevented vascular smooth muscle cell (VSMC) loss and reduced intracerebral haemorrhage^46^. The improved BBB integrity following TGF-β inhibition in our iPSC *COL4A1* cSVD model further supports a gain-of-function mechanism. Although HTRA1 treatment reduced *TGFB1, TGFB2* and *TGFBR2* expression, its benefits likely extend to restoring protease-mediated ECM quality control, including clearance of misfolded mutant collagen IV in the ECM.

Previous work, including ours, shows that ECM integrity is essential for BBB function and that its disruption contributes to cSVD^10, 14, 20, 47–49^. However, the specific role of the pBM in cSVD remains poorly defined, partially because astrocyte-derived pBM is challenging to study *in vivo*. Using our iPSC model, we generated astrocyte-derived pBM and directly assessed its impact on BBB function in cSVD, revealing cell-type-specific neurovascular interactions, which could inform future targeted therapeutic strategies.

Astrocyte-secreted soluble factors are essential for establishing and maintaining the BBB^50^. Consistent with this, conditioned medium from control iPSC-astrocytes restored tight junction defects in *COL4A1*^G755R^ iPSC-BMECs, whereas the mutant astrocyte secretome had reduced ability to do so. Notably, improved TJ protein levels and organisation occurred without corresponding increases in TJ gene expression, suggesting limited involvement of transcription-regulating astrocyte factors such as Sonic Hedgehog (Shh) and Glia-derived neurotropic factor (GDNF) in *COL4A1* cSVD. The exacerbated transcriptomic changes in mutant BMECs when exposing to mutant astrocyte secretome further indicate that cell-autonomous collagen IV effects and astrocyte-derived signals act synergistically to worsen BBB damage. These findings highlight the need for multicellular models to capture the full disease spectrum.

### Limitations

Our study has several limitations. We used conventional 2D cultures of iPSC-astrocytes and BMECs, which clarify individual cell functions but do not capture the complex architecture and signalling of human tissue. Future work using 3D systems, such as brain or vascular organoids and organ-on-chip models, will better reflect *in vivo* cellular organisation. IPSC-astrocytes and BMECs also lack full adult maturity and heterogeneity, a known constraint of current iPSC models. Long-term or vascularised organoids may improve this. Additionally, we focused on a single *COL4A1* variant, G755R. Although this variant represents one of the 85–95% cSVD-associated *COL4A1* missense variants that involve a glycine within the Gly–X–Y motif essential for collagen triple helix formation, including multiple patient lines will be important for validating therapies and assessing genotype-phenotype variations. However, the isogenic control used in this study ensured that observed phenotypes were driven by the *COL4A1^G^*^755^*^R^*variant.

## Materials and methods

### iPSC culture

The *COL4A1*^G755R^ iPSC line derived from a cSVD patient, and its isogenic control line used in this study were reported previously^25^. IPSCs were maintained in Essential 8 medium (E8 medium; Thermo Fisher, 15170001) at 37°C with 5% CO_2_ on Matrigel (Corning, CLS356234) coated 6-well plates. The cells were subculture using 0.5 mM UltraPure EDTA solution (Invitrogen, 15575020) at ∼70% confluence and seeded in E8 medium containing 10 μM Rho kinase (ROCK) Inhibitor (Y-27632) (Sigma, SCM075) on 6-well plates. Y-27632 was withdrawn after 24 hours. IPSCs used in this study were between passages 20 to 40.

### Differentiation of iPSCs into brain microvascular endothelial cells

BMEC differentiation from iPSCs was performed using a protocol adapted from a previous report^30^. IPSCs were seeded on Matrigel-coated 6-well plates in E8 medium with 10 μM ROCK inhibitor at ∼10,000 cells/cm², with daily medium changes. After three days, cultures were switched to DeSR1 containing 6 μM CHIR99021 (StemCell Technologies, 72052) (Day 0). DeSR1 consisted of DMEM/F12 (31331, Life Technologies) supplemented with MEM-NEAA (11140, Life Technologies), GlutaMAX (25030024, Life Technologies), and β-mercaptoethanol (31350, Life Technologies). After 24 hours, the medium was replaced with DeSR2 (DeSR1 plus B27) and refreshed daily until Day 6, when cells were transferred to hECSR1 for 48 hours. The hECSR1 is comprised of human endothelial serum-free medium (hESFM; Thermo Fisher Scientific), 20 ng/mL FGF2 (R&D Systems), 10 μM all-trans retinoic acid (Sigma), and 1 × B27. On Day 8, cultures were changed to hECSR2 (hESFM plus B27). On Day 9, cells were dissociated with TrypLE (Thermo Fisher) and replated onto Matrigel-coated plates or 0.4-μm Transwell inserts (Corning). After 20 minutes, cultures were rinsed with hESFM containing 2% platelet-poor plasma-derived serum and 10 μM Y27632, as a selection step to remove undifferentiated cells, then maintained overnight for attachment.

### Astrocyte differentiation from iPSCs

#### Neuronal progenitor cell (NPC) induction

The iPSCs were first differentiated into NPCs following established protocols^51^. IPSCs were passaged onto Matrigel-coated 6-well plates and cultured in E8 medium supplemented with 10 μM ROCK inhibitor at 100% confluency. After 24 hours, the medium was replaced with neural induction medium (NIM; Day 0), consisting of a 1:1 mixture of DMEM/F12 and Neurobasal supplemented with 1 × B-27, N-2 (Life Technologies, 17502048), 5 μg/ml insulin (Sigma, I9278), 100 μM β-mercaptoethanol, 100 μM nonessential amino acids, 1 mM L-glutamine, 500 μM sodium pyruvate (Sigma, S8636), 50 μg/ml penicillin–streptomycin (Life Technologies, 15140), 10 μM SB431542 (Tocris, 1614), and 1 μM Dorsomorphin (Tocris, 3093). The NIM was refreshed daily. On Day 12, cells were dissociated by 200 ul Dispase (Stem Cell Technologies, 07923), mixed with 2 ml NIM medium, and seeded onto Laminin-coated plates. After an overnight culture (Day 13), NIM was switched to neural maintenance medium (NMM; NIM without SB431542 and Dorsomorphin) supplemented with 20 ng/ml FGF2. On Day 17 when rosettes became visible, cells were dissociated using Dispase and replated onto Laminin-coated plates at a 1:2 ratio. NMM was changed daily. On Day 25, cells were dissociated into single cells using Accutase and replated at a 1:1 ratio on Laminin-coated plates. Medium was refreshed after 24 hours and then every 48 hours. Cells were subsequently passaged at a 1:2 ratio upon reaching 90–100% confluency.

#### Astrocyte differentiation

NPCs were dissociated with Accutase and seeded onto Matrigel-coated 6-well plates in NMM (Day 0). On day 1, medium was replaced with 2 ml STEMdiff^TM^ Astrocyte Differentiation Medium (ADM; Stem cell Technologies, 100-0013), refreshed daily. On Day 7, cells were passaged on Day 7 using Accutase and replated at 1.5-2×10^5^ cells/cm^2^ in ADM, which was then changed every 2-3 days. Additional passages were performed on Day 14 and 21, after which cultures were switched to STEMdiff^TM^ Astrocyte Maturation Medium (AMM; Stem cell Technologies, 100-0016) that was refreshed every 2-3 days. On both Day 28 and Day 35, cells were passaged with Accutase and replated onto Matrigel-coated coverslips in 24-well plates or 6-well plates at 1.5-2×10^5^ cells/cm^2^. After two passages in AMM, astrocyte identity was assessed by immunostaining and RT-qPCR for astrocyte-specific markers, S100β and GFAP.

### Immunofluorescence staining

Cells were washed with PBS and fixed in 4% PFA for 15 minutes at room temperature, then permeabilized with 0.1% Triton X-100. After PBS washes, cells were blocked in 10% donkey serum in PBS for 30 minutes and incubated with primary antibodies in blocking buffer for 1 hour at room temperature (RT), followed by fluorescent secondary antibodies at RT for 1 hour in dark. Antibody details and dilutions are listed in **Table S1 and S2**. Nuclei were counterstained with DAPI, and images acquired using the EVOS™ FLoid™ Cell Imaging Station and analyzed using EVOS and Fiji ImageJ.

### Quantification of TJ Integrity

Two complimentary image analysis approaches were applied. The first measures discontinued membrane staining of TJ markers (ZO-1, Claudin-5, and Occludin), which is presented as numbers of TJ breaks per cell. The second assessed the discontinuous fusion points by measuring the loss of fluorescence intensity of TJ staining at cell junctions, which was presented as discontinued fusion points per cell. Nine images were used, with 3 randomly selected images taken from each experiment, repeated across 3 biological replicates.

### Trans-endothelial electrical resistance (TEER) assay

On day 9 of the iPSC BMEC differentiation, cells were seeded onto 12-well 0.4 μm Trans-well inserts. Twenty-four hours after, trans-endothelial electrical resistance (TEER) measurements were recorded daily using an EVOM2TM Epithelial Volt/Ohm Meter with STX2 electrodes (World Precision Instrument). TEER values were normalized by subtracting the background TEER from a blank well and then multiplied by the surface area (1.12 cm^2^ of 12-well plate) of the Trans-well filter. Results are presented as ohms × cm^2^. All TEER experiments were performed with at least 3 triplicate wells.

### Quantitative real time polymerase chain reaction (RT-qPCR)

Total RNA was extracted using the RNeasy Mini Kit, and 250 ng RNA was reverse-transcribed with the High-Capacity RNA-to-cDNA Kit (Qiagen, 74104). RT-qPCR was performed using PowerUp SYBR Green Master Mix (Thermo Fisher, A25742) on a QuantStudio 3 system (Applied Biosystems) with cycling conditions: 95 °C 10 minutes; 40 cycles of 95 °C 15 seconds, 60 °C 1 minute. The melt curve analysis was performed to confirm amplification specificity (95°C for 15 seconds, 60°C for 60 seconds, then gradual increase to 95 °C). Gene expression was calculated using the 2^−ΔΔCt method with GAPDH as control. Primers are listed in **Table S3**.

### Calcium imaging

IPSC-astrocytes were seeded onto Matrigel-coated four-compartment glass-bottom chamber slides (Cellvis, D35C4-20-1.5-N) at a density of 1 x 10^5^ cells per chamber cultured for 2–3 days. Fluo-4 AM (Invitrogen, ThermoFischer Scientific) was added to each chamber (final concentration: ∼0.5 μM) and incubated for 30 minutes at 37°C in the dark, followed by PBS washes and replacement with fresh astrocyte medium. Calcium activity was stimulated by adding 25 μM L-glutamate (Abcam, ab83389) during live imaging. Fluorescence was recorded on a Zeiss spinning-disk confocal microscope (Axio Observer Z1) using a 20×/0.8 NA objective with 488 nm excitation. Time-lapse images were collected every 0.39 seconds for 30 minutes. Fluorescence traces of individual astrocytes were quantified using ZEN (Zeiss), Fiji (ImageJ), and GraphPad Prism. All experiments were conducted in three independent replicates.

### Glutamate uptake assay

IPSC-astrocytes were seeded in Matrigel-coated 48-well plates at 1 × 10⁶ cells/well and cultured overnight. The next day, medium was replaced with 200 µL pre-warmed serum-free Hank’s Balanced Salt Solution (HBSS; Gibco, 14025092) containing 100 nM glutamate, and cells were incubated for 2 hours at 37 °C. Medium was collected, and cells were washed with cold PBS and lysed in 100 µL cold PBS. Lysates were homogenized by pipetting and centrifuged at 12,000 × g for 15 minutes at 4 °C, and the supernatant was collected. Glutamate levels were measured by mixing 50 µL sample with 50 µL Glutamate Detection Reagent (Glutamate Assay Kit; Abcam, ab83389) and reading luminescence after 60 minutes.

### ECM synthesis and cell seeding

IPSC-astrocytes were seeded onto Laminin (Sigma, L2020) coated 24-well plate at 1–2 × 10⁵ cells/well and cultured for 24 hours before switching to astrocyte medium (AM; Sciencell, SC-1801) supplemented with 50 µg/mL L-ascorbic acid. Medium was refreshed every 24-48 hours for 7 days. The culture was decellularised by incubating with 0.1% Triton X-100 and 20 mM NH₄OH for 5 minutes at RT, followed by gentle rinsing three times (30 seconds each time) with 500 μL PBS per well to remove cell debris. Wells were then incubated overnight at 4 °C with PBS on a gentle shaker and washed again the next day. The resulting ECM was either fixed with 4% PFA for immunostaining or prepared for cell reseeding. Before cell seeding, ECM-coated wells were equilibrated with 1 mL pre-warmed medium for 1 hour at 37 °C. iPSC-BMECs were dissociated with TrypLE and seeded onto the astrocyte ECM at 1 × 10⁴ cells/well in hECSR2 medium containing 10 µM Y-27632. Cultures were maintained at 37 °C and 5% CO₂ with medium changes every 24-48 hours. After 4 days, cells were fixed with 4% PFA for downstream immunocytochemistry.

### Western Blotting

Cells were washed with cold PBS and lysed in RIPA buffer (150 mM NaCl, 1% NP-40, 0.5% sodium deoxycholate, 0.1% SDS, 50 mM Tris HCl, pH 7.4) containing protease and phosphatase inhibitors (Thermo Fisher Scientific) by incubating on ice for 30 minutes with periodic vortexing. Lysates were centrifuged at 12,000 × g for 15 minutes at 4 °C, and the supernatant was collected. The protein samples (20–30 µg) were subjected to 12% SDS-PAGE followed by incubating with primary antibodies in 5% Marvel TBS-T (TBS with 0.1% Tween 20) overnight at 4 °C and then HRP-conjugated secondary antibodies (1:2000 1:5000 dilution) in 5% Marvel TBS-T for 1.5 hours at RT. ECL reagents (Bio-Rad, 1705061) were used to visualise the bands. Images were quantified using Fiji (ImageJ). Antibodies used were listed in **Table S4**.

### HTRA1 and small molecule treatments

Recombinant human HTRA1 protein (20 ng/mL; Abcam, ab134441) or TGF-β inhibitor, SB431542 (10 μM; Cayman Chemical, 1614), were incubated with iPSC-BMECs in hECSR2 medium for 4 days with media refreshed every 48 hours. The cells were then subjected to downstream analysis.

### RNA Sequencing Bioinformatic Analysis

Total RNA was extracted from iPSC-BMECs and astrocytes using the RNeasy Mini Kit (Qiagen, 74104). RNA concentration and purity were measured with a SimpliNano spectrophotometer (Biochrom, 29061711), and samples were submitted to Azenta Life Sciences for sequencing. RNA integrity was verified using an Agilent Bioanalyzer, and only high-quality samples (RIN > 8) were processed further. Paired-end sequencing was performed on an Illumina platform to ensure sufficient transcriptome coverage. Initial raw data processing and alignment to the human reference genome (GRCh38/hg38) were performed by Azenta using a standardized pipeline. Differential expression analysis was performed using DESeq2 (v1.36.0) in R Studio, and significantly differentially expressed genes (DEGs) were defined as adjusted p < 0.05 and |log₂FC| > 1. Principal component analysis (PCA) and hierarchical clustering were used to assesses sample variability. Volcano plots, gene ontology (GO) enrichment, heatmaps, Kyoto Encyclopedia of Genes and Genomes (KEGG) pathway analysis, gene set enrichment analysis (GSEA), and Venn diagrams of differentially expressed genes (DEGs) were performed using the WeiShengXin online platform (https://www.bioinformatics.com). Protein interaction (PPI) networks were constructed with STRING.

### Statistics

All statistical analyses were performed in GraphPad Prism 9. Data normality was assessed using the Shapiro–Wilk test and variance homogeneity was verified with the Brown–Forsythe test. Two-tailed unpaired Student’s t-tests were used for two-group comparisons, and one-way ANOVA with Tukey’s post hoc test for analyses involving more than two groups. Two-way ANOVA with Šídák’s post hoc test was applied when multiple variables were tested. Data are reported as mean ± SEM. The number of biological replicates (n) is specified in each figure legend. For iPSC-derived cells, each n reflects an independent iPSC differentiation unless otherwise noted. Statistical significance was defined as p < 0.05.

## Acknowledgments

The Bioimaging Facility microscopes used in this study were purchased with grants from BBSRC, Wellcome and the University of Manchester Strategic Fund. Special thanks go to Steven Marsden and David Spiller for their help with the microscopy.

