## Supplementary material for "HTRA1 deficiency in *COL4A1* mutant hiPSC-derived astrocytes: a convergent mechanism of cerebral small vessel disease": Supplimental materials

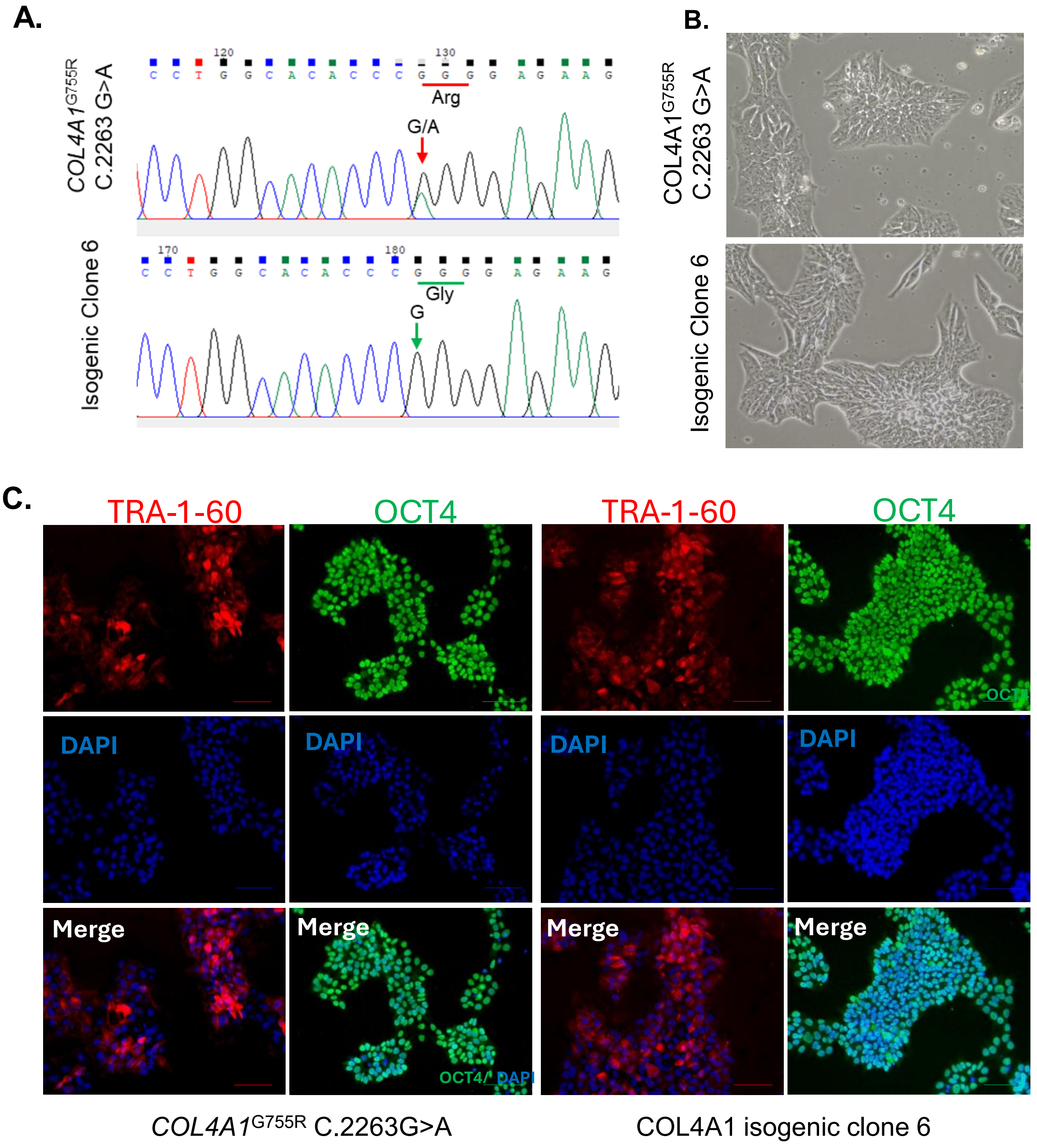


**Figure S1. Characterisation of iPSC lines used in the study.** The iPSC line was generated from a cerebral small vessel disease patient carrying *COL4A1*$^{G755R}$ variant that was corrected using CRISPR/Cas9 gene editing as described in previous publication (ref). (**A**). Sanger sequencing confirmed the existence *COL4A1*$^{G755R}$ variant (top panel) and the correction of the variant (bottom panel). (**B**). Bright field microscopy showing the morphology of the iPSCs. (C). Immunofluorescent staining of pluripotency markers TRA-1-60 and OCT4 in the *COL4A1*$^{G755R}$ and isogenic control iPSC lines. Scale bar, 100 µm.

A.

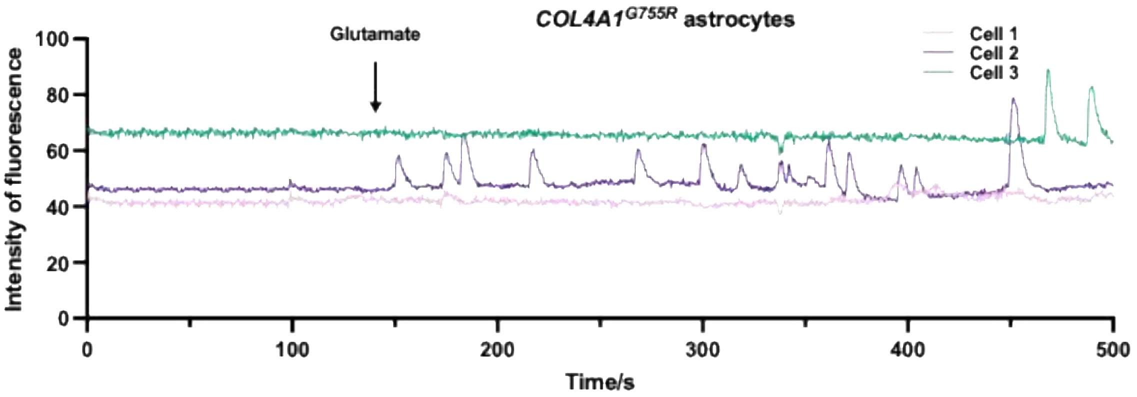


B.

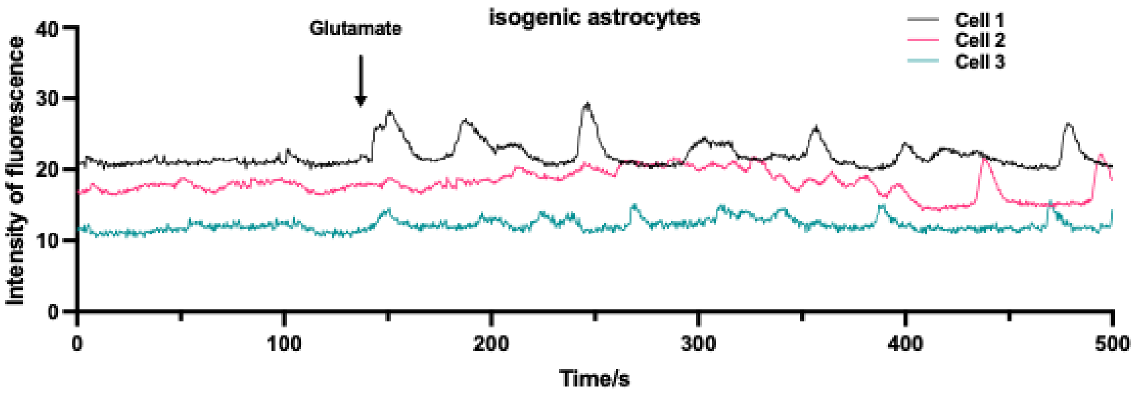


**Figure S2. Calcium imaging.** Astrocytes derived from *COL4A1*$^{G755R}$ (**A**) and isogenic control (**B**) iPSCs were subjected to calcium imaging using Fluo-4 AM indicator. Calcium oscillations waves are shown baseline and upon glutamate stimulation over 500s for 3 selected cells.

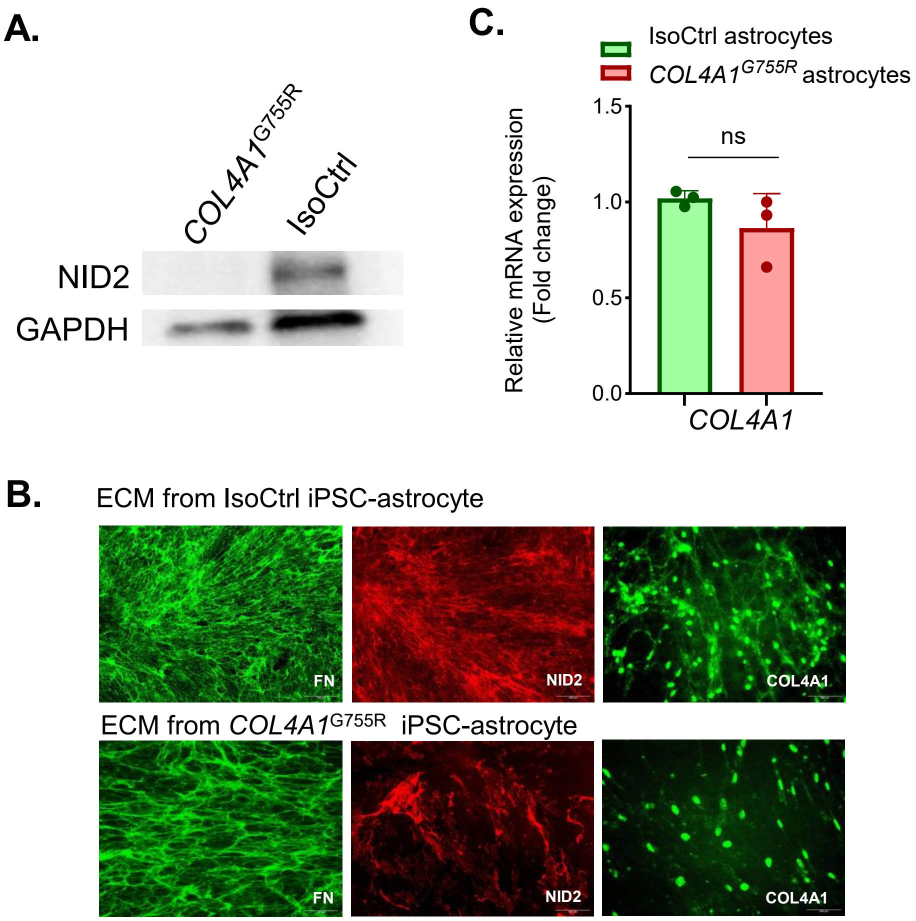


**Figure S3. Extracellular matrix proteins in iPSC-derived astrocytes.** Astrocytes were differentiated from *COL4A1*$^{G755R}$ and isogenic control iPSCs. IPSC-derived astrocytes were subjected to western blotting (**A**) and RT-qPCR (**C**). (**B**). Extracellular matrix (ECM) was prepared from *COL4A1*$^{G755R}$ and isogenic control iPSCs derived astrocytes and immunostained for fibronectin (FN), nidogen 2 (NID2), and Collagen IV $\alpha$1 (COL4A1). Scale bar, 100 µm.

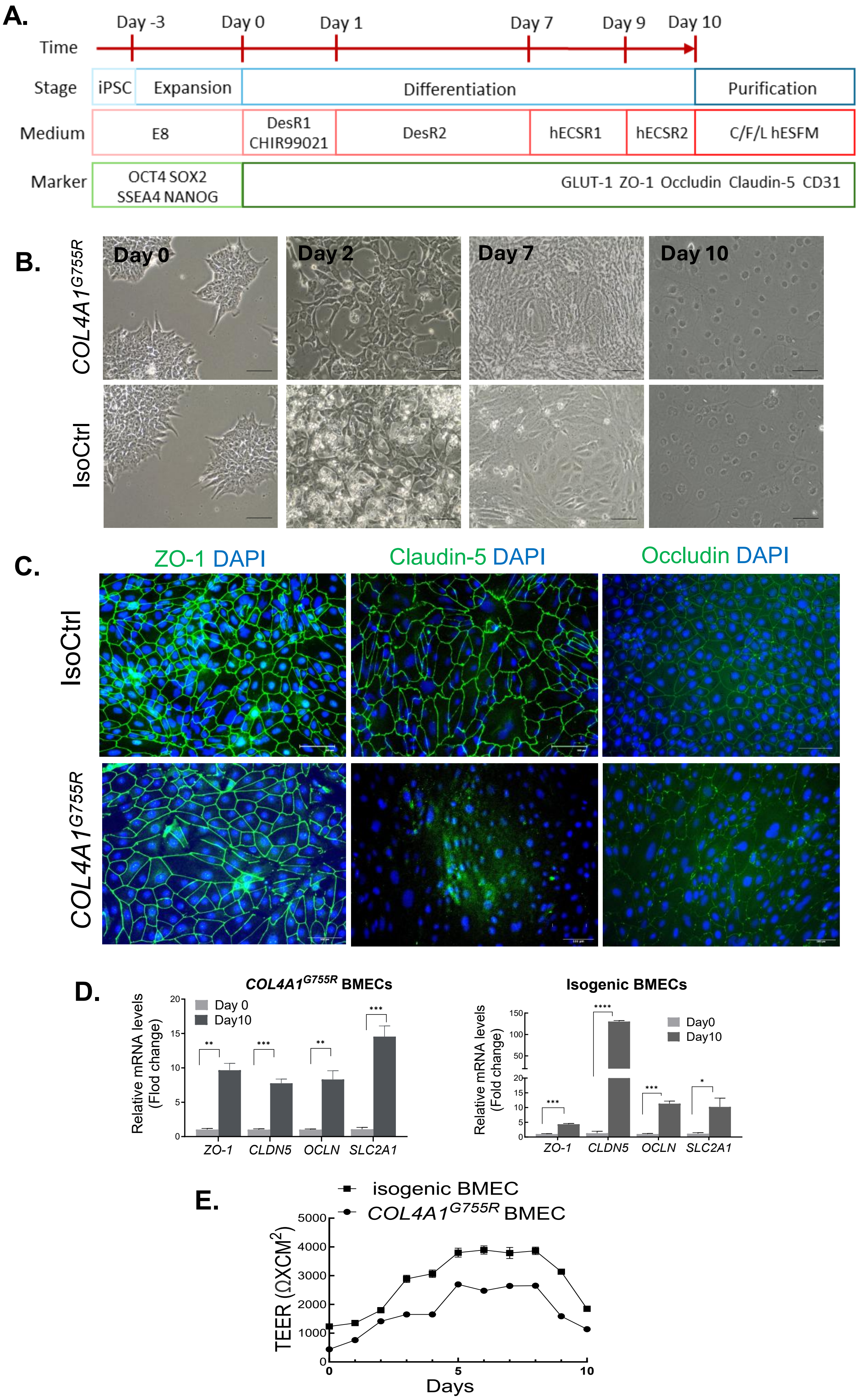
A.
Time
Day -3
Day 0
Day 1
Day 7
Day 9
Day 10
Stage
iPSC
Expansion
Differentiation
Purification
Medium
E8
DesR1 CHIR99021
DesR2
hECSR1
hECSR2
C/F/L hESFM
Marker
OCT4 SOX2 SSEA4 NANOG
GLUT-1 ZO-1 Occludin Claudin-5 CD31
B.
COL4A1G755R
IsoCtrl
Day 0
Day 2
Day 7
Day 10
C.
ZO-1 DAPI
Claudin-5 DAPI
Occludin DAPI
IsoCtrl
COL4A1G755R
D.
COL4A1G755R BMECs
Relative mRNA levels (Flod change)
Day 0
Day10
ZO-1
CLDN5
OCLN
SLC2A1
Isogenic BMECs
Relative mRNA levels (Fold change)
Day0
Day10
ZO-1
CLDN5
OCLN
SLC2A1
E.
isogenic BMEC
COL4A1G755R BMEC
TEER (ΩXCM2)
Days

**Figure S4. BMEC differentiation from *COL4A1*$^{G755R}$ and isogenic control iPSCs and blood-brain barrier (BBB) function.** (**A**). Schematic protocol of BMEC differentiation from iPSCs. (**B**). Phase contrast microscopy showing cell morphological changes during BMEC differentiation. (**C**). Immunofluorescent staining of tight junction (TJ) markers ZO-1, Claudin and Occludin in iPSC-derived BMECs. (**D**). RT-qPCR measurement of expressions of TJ marker genes *ZO-1, CLDN5, OCLD* and *SLC2A1* in *COL4A1*$^{G755R}$ BMECs (left panel) and isogenic control BMECs (right panel). (**E**). Trans-endothelial electric resistance (TEER) measurement of iPSC-derived BMECs. Data in (D and E) are mean ± SEM from triplicate reactions of 3 biological replicates, n=3. Unpaired Student's t-test, *p<0.05, **p<0.01, ***p<0.001, ****p<0.0001. Scale bar, 100 μm.

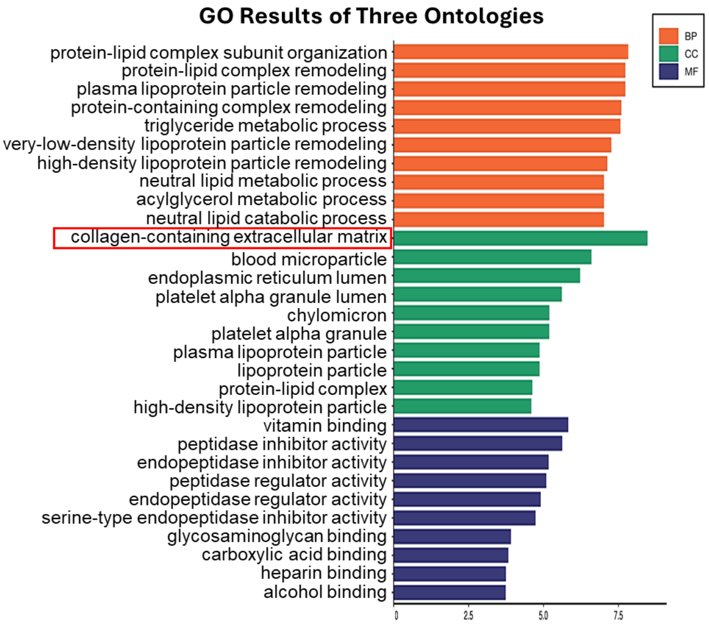


**Figure S5. Gene ontology (GO) analysis of RNA-seq data of iPSC-derived BMECs.** BMECs were differentiated from iPSCs of *COL4A1*$^{G755R}$ and its isogenic control (IsoCtrl) from 3 independent experiments and then subjected to RNA sequencing (RNA-seq). Figure shows GO analysis of differentially expressed genes (DEGs) between *COL4A1*$^{G755R}$ BMECs and IsoCtrol BMECs.

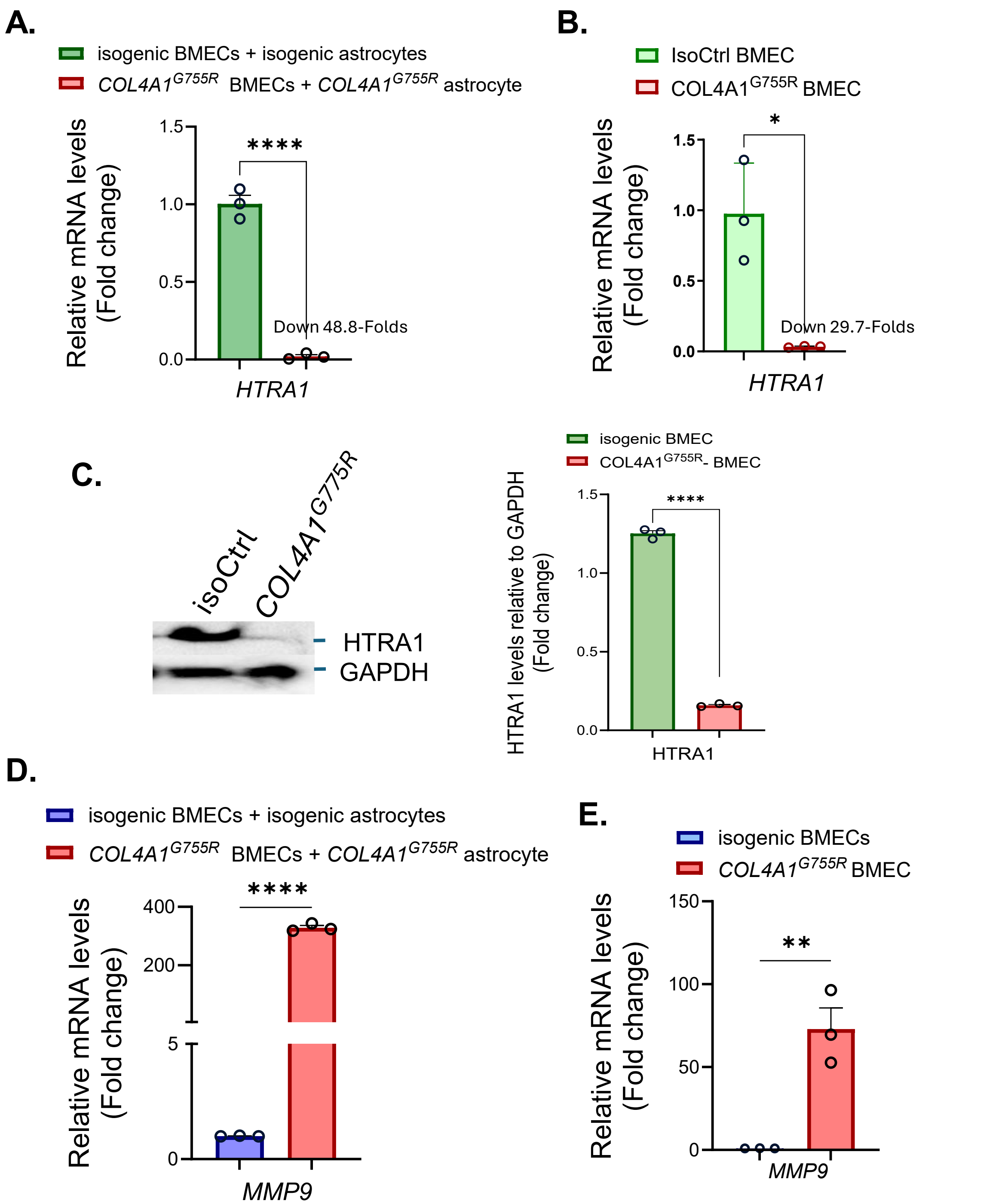


**Figure S6. HTRA1 and MMP9 expression in iPSC-derived BMECs.** (**A**). RT-qPCR determination of HTRA1 mRNA in BMECs derived from *COL4A1*$^{G755R}$ and isogenic control (IsoCtrl) iPSCs. (**B**). RT-qPCR determination of HTRA1 mRNA in BMECs derived from *COL4A1*$^{G755R}$ and IsoCtrl iPSCs which were exposed to their corresponding astrocyte conditioned medium for 4 days. (**C**) Western blotting (WB) of HTRA1 in BMECs derived from *COL4A1*$^{G755R}$ IsoCtrl iPSCs, and its quantification (right panel). (**D**). BMECs derived from *COL4A1*$^{G755R}$ and IsoCtrl iPSCs were exposed to their corresponding astrocyte conditioned medium for 4 days. *MMP9* expression was determined by RT-qPCR. (**E**). RT-qPCR determination of *MMP9* expression in *COL4A1*$^{G755R}$ and IsoCtrl iPSCs derived BMECs. Data in are mean ± SEM from triplicate reactions of 3 biological replicates, n=3. Unpaired Student's t-test, *p<0.05, **p<0.01, ***p<0.001, ****p<0.0001.

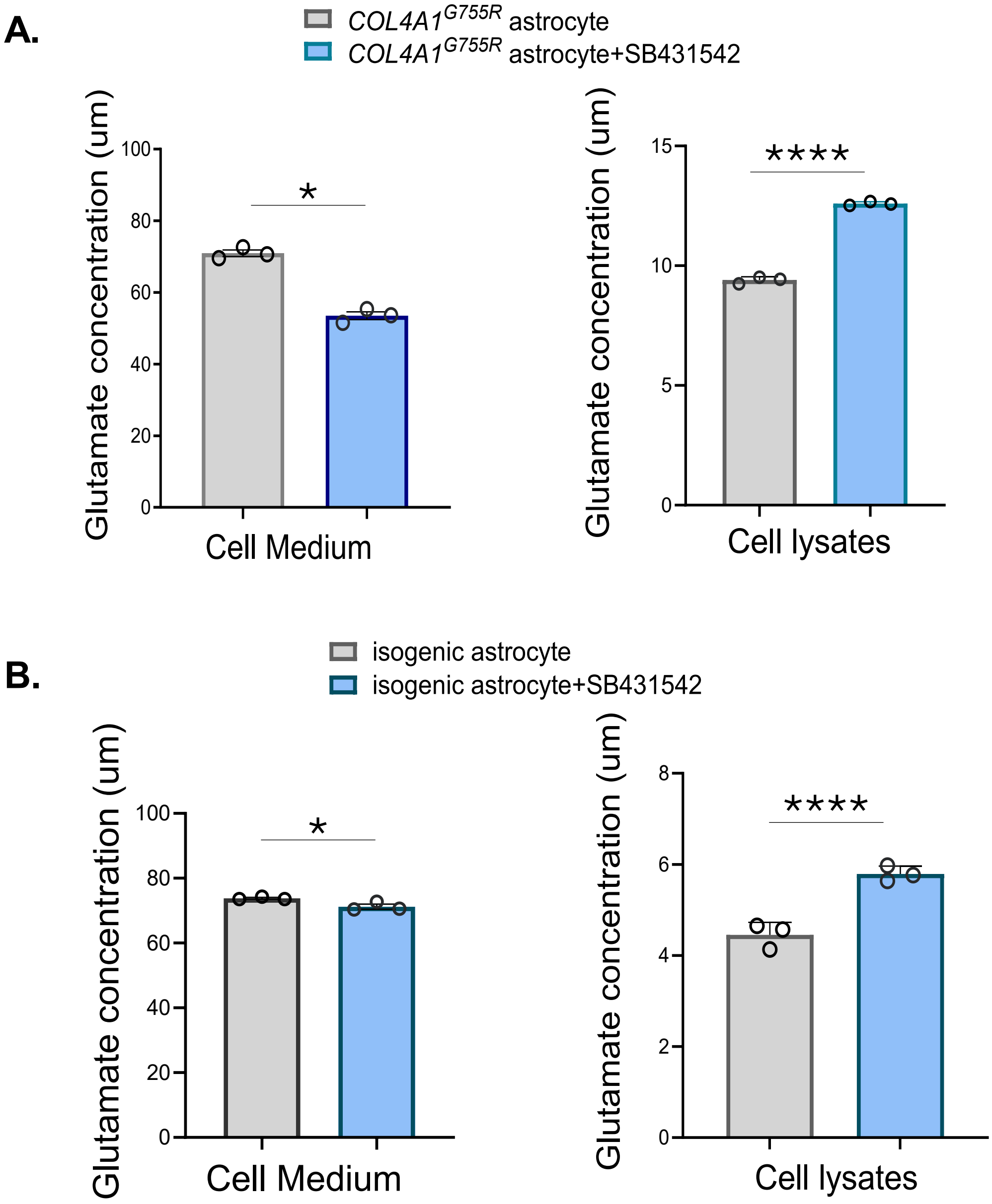


**Figure S7. Inhibition of TGF-β signalling on Glutamate uptake by astrocytes.** Astrocytes derived from *COL4A1*$^{G755R}$ iPSCs (**A**) or isogenic control (IsoCtrl) iPSCs (**B**) were incubated with 100 nM glutamate at 37°C for two hours with or without TGF-β signalling inhibitor SB431542 for 4 days, with medium refreshed every 48 hours. The remaining glutamate in the cell culture medium and cell lysates were measured. Data are means ± SEM from triplicate reactions of 3 biological experiments, n=3. Unpaired Student's *t*-test, *p<0.05. ****p<0.0001.

**Table S1.** Primary antibodies used for immunofluorescent staining.

| Antibody | Target Cell type | Dilution | Host | Supplier | Catalogue no |
|---|---|---|---|---|---|
| Anti-OCT4 | iPSC | 1:250 | Rabbit | Abcam | ab19857 |
| Anti-Tra-1-60 | iPSC | 1:250 | Mouse | Abcam | ab16288 |
| Anti-ZO-1 | BMEC | 1:500 | Rabbit | Invitrogen | UL293821 |
| Anti-GLUT-1 | BMEC | 1:250 | Mouse | Invitrogen | VG3027058 |
| Anti-Occludin | BMEC | 1:250 | Rabbit | Abcam | ab31721 |
| Anti-Claudin-5 | BMEC | 1:250 | Mouse | Invitrogen | UI292618 |
| Anti-PAX6 | NPC | 1:250 | Mouse | Invitrogen | MA1-109 |
| Anti-SOX1 | NPC | 1:250 | Rabbit | Abcam | ab87775 |
| Anti-S100β | Astrocyte | 1:250 | Goat | R&D | AF1820 |
| Anti-GFAP | Astrocyte | 1:500 | Rabbit | Invitrogen | UE286801 |
| Anti-Fibronectin | ECM/Astrocyte | 1:500 | Mouse | Abcam | ab281575 |
| Anti-HTRA1 | ECM/Astrocyte | 1:250 | Rabbit | Abcam | ab274322 |
| Anti-NID2 | ECM | 1:250 | Rabbit | Invitrogen | PA5-62615 |
| Anti-p-SMAD2/3 | ECM/Astrocyte | 1:250 | Rabbit | Abcam | ab272332 |
| TGFβ1 | ECM/Astrocyte | 1:250 | Rabbit | Abcam | ab215715 |
| Anti-Collagen | ECM | 1:250 | Mouse | NOVUSBIO | 8361B-021120-AF405 |

**Table S2.** Secondary antibodies used for immunofluorescent staining.

| Antibody | Excitation wavelength | Dilution | Target Species | Supplier | Catalogue no |
|---|---|---|---|---|---|
| Donkey -anti-mouse | 488 | 1:500 | Mouse | Abcam | ab150105 |
| Donkey-anti-mouse | 594 | 1:500 | Mouse | Abcam | ab150108 |
| Donkey -anti-rabbit | 488 | 1:500 | Rabbit | Abcam | ab150073 |
| Donkey -anti-rabbit | 594 | 1:500 | Rabbit | Abcam | ab150076 |

**Table S3.** Primers used for RT-qPCR.

| *Target Gene* | **Forward sequence (5'-3')** | **Reverse sequence (5'-3')** |
|---|---|---|
| ***GAPDH*** | CATGTTCGTCATGGGTGTGAACCA | ATGGCATGGACTGTGGTCATGAGT |
| ***Zo-1*** | ACCAGTAAGTCGTCCTGATCC | TCGGCCAAATCTTCTCACTCC |
| ***Claudin-5*** | GTTCGCCAACATTGTCGTCC | GTAGTTCTTCTTGTCGTAGTCGC |
| ***Occludin*** | GACTTCAGGCAGCCTCGTTAC | GCCAGTTGTGTAGTCTGTCTCA |
| ***GLUT-1*** | AGGTGATCGAGGAGTTCTAC | TCAAAGGACTTGCCCAGTTT |
| ***PAX6*** | GTGTCCAACGGATGTGTGAG | CTAGCCAGGTTGCGAAGAAC |
| ***SOX1*** | CCTCCGTCCATCCTCTG | AAAGCATCAAACAACCTCAAG |
| ***S100β*** | TGTAGACCCTAACCCGGAGG | TGCATGGATGAGGAACGCAT |
| ***GFAP*** | GTCCCCCACCTAGTTTGCAG | TAGTCGTTGGCTTCGTGCTT |
| ***HTRA1*** | TCCCAACAGTTTGCGCCATAA | CCGGCACCTCTCGTTTAGAAA |
| ***NID2*** | AAGCCCGATTCAGCAACCTC | CACATACTGCGTTTCCCTGGG |
| ***FBLN5*** | CTCACTGTTACCATTCTGGCTC | GACTGGCGATCCAGGTCAAAG |
| ***FBLN2*** | CAGGTGGCCTCTAACACCATC | CTGCTTGCAGGGTCCATTGT |
| ***FBN1*** | TTTAGCGTCCTACACGAGCC | CCATCCAGGGCAACAGTAAGC |
| ***ECM2*** | CCGAATGCCCTCTCGATCC | TGGGTAAGCATGGCGTTGATG |
| ***ADAMTS2*** | GACACGGGCCACGATGAATA | GGTGACAGGAGCATAGCCTT |
| ***LTBP1*** | GCTTCCGTCCAGATACATCAG | CTTGGTACGAGACTTGGGATTG |
| ***TGFβ1*** | GGCCAGATCCTGTCCAAGC | GTGGGTTTCCACCATTAGCAC |
| ***TGFβ2*** | CAGCACACTCGATATGGACCA | CCTCGGGCTCAGGATAGTCT |
| ***TGFBR2*** | GTAGCTCTGATGAGTGCAATGAC | CAGATATGGCAACTCCCAGTG |
| ***PDGFβ*** | CTCGATCCGCTCCTTTGATGA | CGTTGGTGCGGTCTATGAG |
| ***ICAM5*** | TCCCCGGCTCTTGGAAGTT | CAGGACTCAGATTCTGGTCCC |
| ***KDR*** | GGCCCAATAATCAGAGTGGCA | CCAGTGTCATTTCCGATCACTTT |
| **SNAI1** | ACTGCAACAAGGAATACCTCAG | GCACTGGTACTTCTTGACATCTG |

**Table S4.** Antibodies used for western blotting.

| Antibody | Dilution | Host | Supplier | Catalogue no |
|---|---|---|---|---|
| Anti-GAPDH | 1:1000 | Rabbit | Abcam | ab9485 |
| Anti-HTRA1 | 1:1000 | Rabbit | Abcam | ab274322 |
| Anti-ZO-1 | 1:500 | Rabbit | Invitrogen | UL293821 |
| Anti-GLUT-1 | 1:250 | Mouse | Invitrogen | VG3027058 |
| Anti-Occludin | 1:250 | Rabbit | Abcam | ab31721 |
| Anti-Claudin-5 | 1:250 | Mouse | Invitrogen | UI292618 |
| Anti-NID2 | 1:250 | Rabbit | Invitrogen | PA5-62615 |
